# Universal functions of prion candidates across all three domains of life suggest a primeval role of protein self-templating

**DOI:** 10.1101/2022.05.30.493841

**Authors:** Tomasz Zajkowski, Michael D. Lee, Siddhant Sharma, Alec Vallota-Eastman, Mikołaj Kuska, Małgorzata Malczewska, Lynn J. Rothschild

## Abstract

Amyloid-based prions have simple structures, a wide phylogenetic distribution, and a plethora of functions in contemporary organisms, suggesting they may be an ancient phenomenon. However, this hypothesis has yet to be addressed with a systematic, computational, and experimental approach. Here we present a framework to help guide future experimental verification of candidate prions with conserved functions in order to understand their role in the early stages of evolution and potentially in the origins of life. We identified candidate prions in all high-quality proteomes available in UniProt computationally, assessed their phylogenomic distributions, and analyzed candidate-prion functional annotations. Of the 27,980,560 proteins scanned, 228,561 were identified as candidate prions (∼0.82%). Among these candidates, there were 84 Gene Ontology (GO) terms conserved across the 3 domains of life. We found that candidate prions with a possible role in adaptation were particularly well-represented within this group. We discuss unifying features of candidate prions to elucidate the primeval roles of prions and their associated functions. Candidate prions annotated as transcription factors, DNA binding, and kinases are particularly well suited to generating diverse responses to changes in their environment and could allow for adaptation and population expansion into more diverse environments. We hypothesized that these functions could be evolutionarily ancient, even if individual prion domains themselves are not evolutionarily conserved. Candidate prions annotated with these universally-occurring functions potentially represent the oldest extant prions on Earth and are therefore excellent experimental targets.

## 1. Introduction

### 1.1 Amyloids and prions

Many proteins can form fibrillar aggregates called amyloids. These were originally discovered as factors involved in the development of neurodegenerative diseases such as Alzheimer’s, Parkinson’s, and Huntington’s diseases, but have since been shown to have a role in many nonpathogenic and, indeed, beneficial physiological functions as well (Chiti and Dobson 2006; Hafner Bratkovič 2017; Inge-Vechtomov, Zhouravleva, and Chernoff 2007). Three lines of argumentation suggest that amyloids might have been present on Earth even before the first forms of what we would consider life evolved. First, prebiotically plausible peptides can form amyloid fibrils (Gazit 2007; Rufo et al. 2014). Second, the amyloid fold (the structure when proteins form amyloid fibrils) is considered the lowest point on the energy landscape for protein folding and aggregation, therefore amyloidogenesis is considered as an intrinsic property of polypeptides (Jahn and Radford 2005; Dobson 2003). Third, amyloids bear a phenomenological similarity to crystals and could have occurred abiotically (Yoshimura et al. 2012). Given these arguments, it is expected that amyloids should be old and very common in nature. Supporting this point of view, amyloids are known to perform multiple physiological functions in diverse organisms from microorganisms to humans. Organisms capitalize both on the physical rigidity of pre-formed fibrils as well as on the process of aggregation itself. For example, the durability and strength of amyloid fibers are conveyed in the properties of silkworm and spider silk (Slotta et al. 2007; Shi et al. 2014). In humans, functional amyloids build scaffolds on which melanin (skin pigment) is deposited (Bissig, Rochin, and van Niel 2016). In bacteria and fungi, highly-conserved amyloid-forming proteins are components of the extracellular matrix and contribute to cell adhesion and biofilm formation (Beyersdörfer 2019; Garcia et al. 2011).

Other physiological processes rely on the ability of a protein to exist in two different states: soluble (an individual protein, capable of carrying out its typically understood function(s)) and aggregated (multiple proteins aggregated into the amyloid form, not capable of carrying out their typically understood individual function(s)). The amyloid formation has characteristics of autocatalysis and after reaching a certain threshold of aggregation often leads to the quick depletion of the soluble fraction of protein that becomes trapped in the form of fibrillar aggregates (Sabaté, Gallardo, and Estelrich 2003). This ability to exist in two very different and self-excluding states (soluble or aggregated) allows some amyloid-forming proteins to function as molecular switches (Brown and Lindquist 2009; Jarosz and Khurana 2017; Halfmann and Lindquist 2010). In some cases, the amyloid form can be inherited cytoplasmically during cellular replication, where it will continue the conversion of a soluble fraction of freshly synthesized protein into the non-functional aggregated fibrillar form. A cell harbouring such self-templating and self-perpetuating aggregation of a protein phenotypically resembles a cell with a deletion of the gene that codes for this protein (Wickner, Edskes, and Shewmaker 2006). These heritable protein aggregates are called *prions*. While some proteins can be prions without being amyloid-based, like intrinsically disordered RNA-binding proteins in *S. cerevisiae* (Chakrabortee, Byers, et al. 2016), here we focus on amyloid-based prions.

Prions can be sustained for many generations (Krammer, Schätzl, and Vorberg 2009). An individual protein that contributes to the prion aggregate is called a “prion protein”, and a distinct protein domain that is responsible for prion aggregation is called a prion domain (PrD). Just like amyloids, prions were first discovered in association with diseases but are now known to regulate a variety of beneficial physiological functions. For example, in plants, prions regulate flowering time (Chakrabortee, Kayatekin, et al. 2016). In yeast, among numerous other functions, prions can drive large shifts in metabolism, regulating whether cells are metabolic specialists or generalists (Brown and Lindquist 2009). In bacteria, prion aggregation can regulate translation (Pallarès and Ventura 2017) and plasmid copy-number (Molina-García et al. 2016). Prion inheritance is often referred to as epigenetic; epi-(□πι-“over, outside of, around”) implies features that are “on top of” the traditional genetic basis for inheritance. Some prions even regulate other mechanisms of epigenetic inheritance via chromatin modification (Harvey et al. 2020; Goncharoff, Du, and Li 2018).

Examples of proteins capable of forming prions are widespread on the phylogenetic tree of life, showing that prion formation is common and that it may be an early-evolved cellular mechanism. It is not clear if this mechanism did indeed evolve early and has been conserved as many forms of life have diverged, or if it has evolved multiple times independently via convergent evolution. We call the second hypothesis *the recurring domestication of prions*.

To pursue the question of the origin of prions, we searched ∼5,000 high-quality proteomes for candidate prion proteins (cPrPs), and then functionally annotated these candidates. We then focused on functional annotations of cPrPs that were identified in all three domains of life. We suspected that these conserved functions associated with cPrPs may be evolutionarily linked to prion aggregation. Determining the functions of the most conserved prion proteins might help unravel the role of the prion phenomenon in biology today, its evolutionary age, and the role it might have played in the origin of life.

### 1.2 Conservation and distribution of prion functions on the tree of life

Different types of prion-forming domains exist. Most known PrDs are enriched in glutamine and asparagine (Q/N enriched) and depleted in charged amino acids. At least one known prion protein, – the mammalian PrP, does not have this bias but is still characterized by an intrinsically disordered domain. In both cases, prion proteins can be conserved across organisms. The first prion-forming protein ever described PrP – has no Q/N bias and is conserved up to early chordates (Holmes et al. 2013; Ehsani et al. 2011; Westaway et al. 2011) which gave rise to early tetrapods about 380 million years ago (Long and Cloutier 2022). The Q/N bias in the prion domain (PrD) of the translation terminator (Sup35) is found in both *Ascomycota* and *Basidiomycota*, which diverged more than 1 billion years ago (L. B. Harrison et al. 2007). While many Q/N-rich prion proteins have been identified as conserved between different types of yeast, no conservation was detected in Q/N-rich prion proteins between humans and yeast (An and Harrison 2016). And so far no conservation has been shown to reach across different domains of life.

Prion proteins can be highly similar in sequence, or more often similar in the chemical properties of the sequence (Su and Harrison 2019). In *S. cerevisiae*, Ure2p and Sup35p sustain their ability to form prions even when the amino-acid sequence of their prion domains is “scrambled,”as long as they maintain the same general chemical properties of the amino-acid composition (Ross, Baxa, and Wickner 2004; Ross et al. 2005). The fact that prion aggregation depends on the composition of prion domains and is (somewhat) independent of exact primary sequence challenges conventional bioinformatics approaches when applied to prions. Nevertheless, Su and Harrison (Su and Harrison 2019) were able to identify prion domains that are both conserved in the primary sequence and amino-acid composition in *S. cerevisiae*. In their dataset, the most conserved prion-like domain across the *Saccharomyces* genus was the protein NRP1, originally described as a prion by Alberti et al. (Alberti et al. 2009). NRP1 is a putative RNA-binding protein that localizes to dense amorphous aggregations in the cytosol called stress granules (Buchan, Muhlrad, and Parker 2008). Other examples of confirmed conserved prions include GLFG-motif nucleoporin NUP100, a confirmed prion-forming protein, which is part of the nuclear pore complex (Halfmann et al., 2012); GLN3, a transcriptional activator of genes regulated by nitrogen catabolite repression; RBS1, a protein involved in the assembly of the RNA polymerase III (Pol III) complex (Cieśla et al. 2015); and MED3/PGD1, a subunit of the RNA polymerase II mediator complex (Bourbon 2008). Having a role in RNA binding and gene regulation is a common theme for many prion candidates, not only in *Saccharomyces* (A. F. Harrison and Shorter 2017; Nizhnikov et al. 2016), and is often connected to phase transitions (Malinovska, Kroschwald, and Alberti 2013), again, promoting the formation of stress granules (Kato et al. 2012; Kroschwald et al. 2015).

Our current information on confirmed prions is limited because it is difficult and time-consuming to experimentally verify them. To broaden our understanding of the ancient functions of prions beyond this small group of confirmed prions, we include functions of proteins that are predicted to behave as prions. Development of prion-prediction algorithms has enabled proteome-wide analysis of proteins revealing thousands of candidate prions (Cascarina and Ross 2020; Espinosa Angarica, Ventura, and Sancho 2013; Michelitsch and Weissman 2000; P. M. Harrison and Gerstein 2003; Alberti et al. 2009; Afsar Minhas, Ross, and Ben-Hur 2017; Sabate et al. 2015; Ross et al. 2013; Lancaster et al. 2014; Zambrano et al. 2015). These efforts were recently reviewed by Batlle and Gil-Garcia (Gil-Garcia et al. 2021; Batlle et al. 2017). The most widely used prediction program is Prion-Like Amino Acid Composition (PLAAC) (Chakrabortee et al. 2016; Lancaster et al. 2014). Despite PLAAC’s underlying machine learning algorithm being trained on yeast PrDs, it has proven successful in facilitating the identification of prions in the domain Archaea (Zajkowski et al. 2021), Bacteria (Yuan and Hochschild 2017; Fleming et al. 2019; Pallarès and Ventura 2017; Molina-García et al. 2016), and even in viruses and phages (Tetz and Tetz 2018; Tetz and Tetz 2017; Nan et al. 2019). This observation alone speaks to the chemically conserved nature of at least a subset of cPrDs across all domains of life.

Several studies have made a rudimentary attempt to describe a phylogenetic distribution of predicted prion candidates in all domains of life. In one such study, Angarica noticed that there are much fewer prion candidates in the domains Bacteria and Archaea than in Eukarya (Espinosa Angarica et al. 2014). This may be because bacterial and archaeal proteomes appear to have fewer Q/N-rich regions in general as compared with eukaryotes (Michelitsch and Weissman 2000). Investigations into the phylogenetic relationships of prion candidates were further augmented with analysis of predicted function derived from Gene Ontology (GO) annotations (Ashburner et al. 2000). Angarica et al. performed the first wide-ranging study of prion candidates in proteomes (Espinosa Angarica, Ventura, and Sancho 2013). Using GO annotations, they observed that predicted prionogenic domains (or candidate prion domains -cPrDs) co-exist with different functional domains of proteins that localize to different cellular compartments, depending on the taxon and organism group. In bacteria, cPrDs are significantly enriched in proteins with annotations involved with the cell wall. Accordingly, bacterial candidate prion proteins (cPrPs) appear to be involved in metabolic and catabolic processes resulting in the construction and disassembly of the cell wall (Espinosa Angarica, Ventura, and Sancho 2013). Another study similarly showed that bacterial prion candidates are associated with peripheral rearrangement, macromolecular assembly, cell adaptability, and invasion (Iglesias, de Groot, and Ventura 2015). This logical convergence of GO cellular localization and function encourages further use of this method as a potentially valuable tool for studying global trends in the roles of prions in biology.

In our previous work, we identified prion candidates in archaea using PLAAC and looked into their GO annotations. Similar to what Angarica et al. found in bacteria, many prion candidates were involved in the construction and adhesion of the cell wall. Archaeal prion candidates were also significantly functionally enriched in regulation through transcription, calcium-binding, and copper ion binding (Zajkowski et al. 2021) – all of which could be evolutionarily early-evolved functions. As one example, the resistance to copper toxicity would have been necessary in early Earth environments (Puig and Thiele 2002). It is, therefore, possible that prion domains may be essential to the function of many proteins involved in copper binding, and may have facilitated the evolution of copper tolerance in early microbial life.

Very recently Garai et al. analyzed the functional divergence of prion candidates in plants (Garai et al. 2021). They found the highest density of prion candidates in green algae. They concluded that the prion phenomenon of aggregation has been conserved from chlorophytes to angiosperms, possibly offering an advantage during the evolution of plants. Similar to bacteria, unicellular algae might have benefited from the physical properties of amyloid-based prions contributing to cell-cell adhesion of their biofilms (Jarvis and Mostaert 2012). Rice also had a high number of candidate prion proteins, with the candidate prions significantly enriched in transposons/retrotransposons. Authors of this analysis suggested the role of candidate prions is in stress response and memory pathways in the plant kingdom (Garai et al. 2021). These observations suggest the role of plant prions in adaptation to changing environments – an adaptation which has been proven in the case of yeast (Oamen et al. 2020; Itakura et al. 2020). It is quite possible that the role of prions in facilitating adaptation could be an evolutionarily ancient one even if individual prion proteins themselves are not evolutionarily conserved.

Putative prion domains rich in Q/N similar to those found in yeast are often detected in large numbers in eukaryotic proteomes. For example, in the genomes of *Drosophila melanogaster, Plasmodium falciparum, Helobdella robusta*, and *Dictyostelium*, more than 20% of proteins contain cPrDs (Malinovska et al. 2015; An and Harrison 2016; Pallarès et al. 2018). GO annotations of prion candidates in *Plasmodium* point to their role in the regulation of gene expression similar to those described by Garai et al. in plants.

Taken together, previous work suggests that the prion phenomenon is widespread but can be randomly distributed even among closely related species. Functions associated with prion proteins vary widely, but functions associated with survival and regulation seem to be more common than others. This tendency can be observed in organisms belonging to different kingdoms within Eukarya as well as across different domains of life, indicating a substantial consistency in the conservation of prion-related functions across different evolutionary epochs. So far, the analysis of prions in proteomes has been mostly focused on selected taxa, and the evolutionary conservation of prion proteins is mostly studied within taxonomic groups. Researchers agree that some prions can be very evolutionarily old, but the question of “how old?” is rarely asked. Identifying a subset of the potentially earliest evolved prions on Earth by looking for similar sequences and functions in a large sampling of all three domains of life could be a valuable step toward understanding the roles of prions in evolution – as well as their roles in contemporary organisms including the pathology of devastating amyloid-associated disorders. To this end, we analyzed all high-quality proteomes available in UniProt using prion-prediction software to prepare a list of candidate prion proteins whose functions are common to all domains. We discuss unifying features of these candidate prions in an effort to elucidate the primeval roles of prions and their associated functions. And we present a framework to help guide future experimental verification of candidate prions with conserved functions in order to understand their role in the early stages of evolution and potentially in the origins of life.

Experimental verification and detailed analysis of these candidates could reveal the possible role(s) of prions at the very early stages of evolution, including the time of the last universal common ancestor (LUCA).

## 2. Methods

### Computational identification of prion candidates

Reference proteomes for Archaea, Bacteria, and Eukarya were downloaded from the UniProt database (UniProt Consortium 2021); accessed 1-Jun-2021) including their stored Gene Ontology (GO) annotations (UniProt Consortium 2021; Gene Ontology Consortium 2021). Proteomes possibly of low quality were excluded by retaining only those ranked as “standard” based on UniProt’s “Complete Proteome Detector” algorithm. This resulted in: 1,151 archaeal proteomes containing 2,182,921 proteins; 2,836 bacterial proteomes holding 10,939,944 proteins; and 925 eukaryal proteomes holding 14,857,695 proteins. The command-line version of PLAAC (Alberti et al. 2009; Lancaster et al. 2014), installed from https://github.com/whitehead/plaac on 9-Sep-2020, was used to identify proteins with candidate prion domains in each proteome individually with default settings other than “-a 0”. Unless otherwise noted, those with a COREscore > 0 (other than “NA”) were considered to contain a candidate prion domain (cPrD).

### GO enrichment analysis

GO enrichment analyses were performed to identify enrichment (i.e., over-representation) or purification (i.e., under-representation) of frequencies of GO annotations in proteins with candidate prion domains as compared to all proteins scanned. This was performed with the goatools v0.8.12 (Klopfenstein et al. 2018) “find_enrichment.py” script with default settings, which includes propagating counts to parents, and was done for each domain of life separately. Statistical significance was defined as those with Benjamini-Hochberg false-discovery rates of ≤ 0.05. Plots of enrichment values were generated with R v3.6.3 (R Core Team. 2020), with the ggplot2 package, v3.3.5 (H. Wickham. 2016). The venn diagram was generated with the R package ggvenn v0.1.9 (Linlin Yan.2022).

### Phylogenomic tree construction

Phylogenomic trees for each domain were produced with GToTree v1.6.11 (Lee 2019), using the prepackaged single-copy gene sets for archaea (76 target genes) and bacteria (74 target genes), and a universal set (16 target genes) for eukarya (Hug et al. 2016). Within GToTree, target genes were identified with HMMER3 v3.2.2 (Eddy 2011), individually aligned with muscle v3.8.1551 (Edgar 2004), trimmed with trimal (Capella-Gutiérrez et al. 2009), and concatenated before phylogenetic estimation with FastTree2 v2.1.10 (Price et al. 2010). The trees were initially visualized and edited through the Interactive Tree of Life site (Letunic and Bork 2007).

### KO annotation of prion candidates

In addition to utilizing the GO annotations that were available with the UniProt data, functional annotation of candidate prion proteins based on Kegg Orthology (KO) terms (Kanehisa et al. 2016) was performed with KOfamScan v1.3.0 (Aramaki et al. 2019).

### Additional data and code availability

All of the UniProt data accessed on 1-June-2021 and utilized in this work along with annotated code and additional data files too large to include here (such as annotation and sequence information for all proteins) is available at https://figshare.com/projects/Zajkowski_et_al_2022_3-domain_prion_data_and_code_repository/133155. Additionally, conda (Anaconda Software Distribution) was utilized for program installation and environment management, and bit v1.8.42 (Lee 2022) and TaxonKit v0.9.0 (Shen and Ren 2021) were utilized heavily for helper scripts and manipulating taxonomy IDs.

## 3. Results

### 3.1 Computational identification and phylogenetic distribution of prion candidates

#### 3.1.1 Archaea

A total of 1,151 archaeal proteomes containing 2,182,921 proteins were scanned with PLAAC (Table S1). Of these, 728 proteomes had at least 1 cPrD detected (∼63%), and 2,191 proteins contained a cPrD (∼0.1%). The frequency of identified cPrDs within each archaeal proteome was ∼1.90 ± 2.78 (mean ± 1 SD), with a median of 1 (Table S2). Normalized per 1,000 proteins per proteome (permil), it was 1.12 ± 1.67, with a median of 0.6. The distribution of those with at least one cPrD detected spans the archaeal proteomes scanned, as depicted by the blue tips in Figure 1. Those with a higher density of cPrDs (meaning when normalized to per 1,000 proteins, “permil”) included members of the classes Methanomicrobia (>= 10 permil) and Methanobacteria and candidatus Micrarchaeota (>= 15 permil; Figure 1; Table S2).

**Figure 1.**
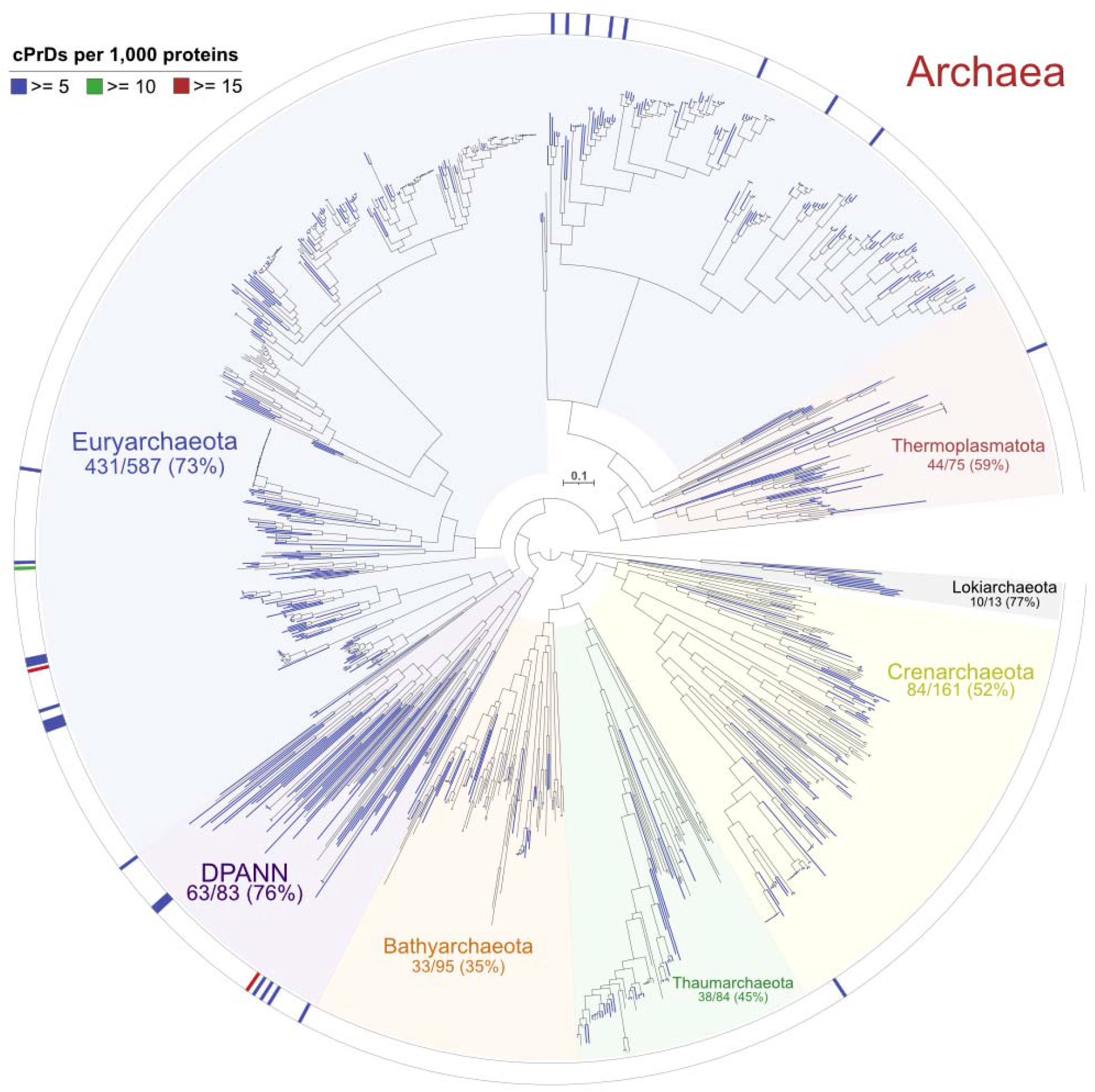
Phylogenomic tree of analyzed archaeal proteomes with their distribution of cPrDs overlain. Tips with at least 1 cPRD identified are colored blue. The outer ring depicts those with greater than 5, 10, or 15 cPrDs per 1,000 proteins within a proteome.

#### 3.1.2 Bacteria

For Bacteria, 2,836 proteomes containing 10,939,944 proteins were scanned (Table S1). Of these, 2,547 proteomes had at least 1 cPrD detected (∼90%), and 15,861 proteins contained a cPrD (∼0.15%). Counts of identified cPrDs in bacterial proteomes averaged ∼5.59 ± 5.97 (mean ± 1SD) with a median of 4. Normalized this was ∼1.39 ± 1.40 with a median of 1.04 per 1,000 proteins (permil). Those with relatively higher permil values included a member of the genus *Candidatus Deianiraea* of the Alphaproteobacteria class (>= 10 permil) and members of the genus *Mycoplasma* within the Phylum Tenericutes (>= 15 permil; Figure 2; Table S2).

**Figure 2.**
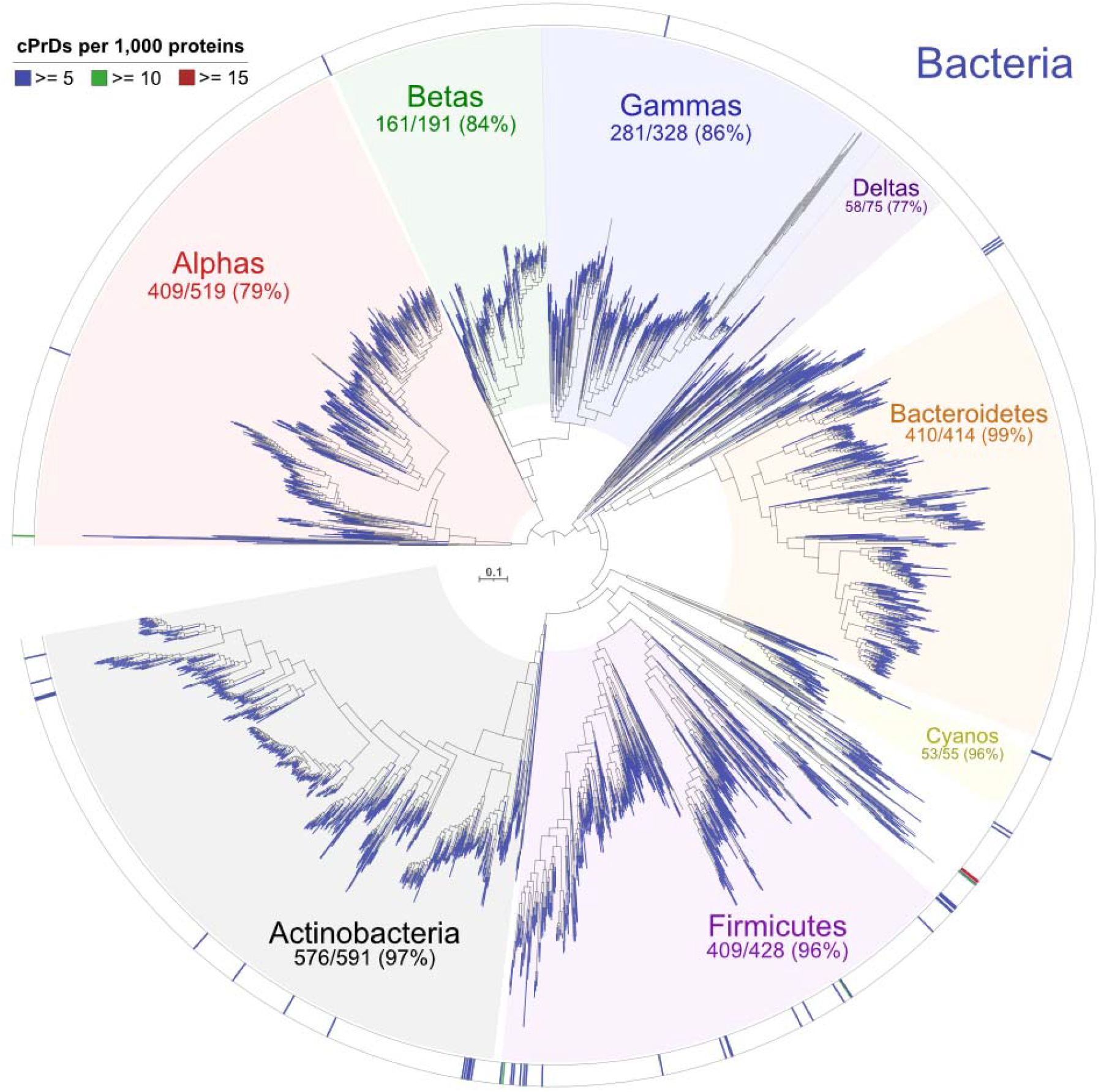
Phylogenomic tree of analyzed bacterial proteomes with their distribution of cPrDs overlain. Tips with at least 1 cPRD identified are colored blue. The outer ring depicts those with greater than 5, 10, or 15 cPrDs per 1,000 proteins within a proteome.

#### 3.1.3 Eukarya

A total of 925 Eukarya proteomes containing 14,857,695 proteins were scanned with PLAAC (Table S1). Of these, 907 proteomes had at least 1 cPrD detected (∼98%), and 210,509 proteins contained a cPrD (∼1.4%). The frequency of cPrDs in eukaryal proteomes was ∼228 ± 154.4 (mean ± 1SD) with a median of 189. Normalized to per 1,000 proteins (permil) this was ∼16.31 ± 12.24, with a median of 12.43 (Table S2). There were generally many more cPrDs identified in the Eukarya domain, but as PLAAC was developed based on known eukaryal prions, it should not currently be concluded this is a true trend and not a methodology bias. One quickly apparent trend in terms of density of cPrDs in Eukarya is that there are consistently lower permil values in the Phylum Chordata (Figure 3; Table S2).

**Figure 3.**
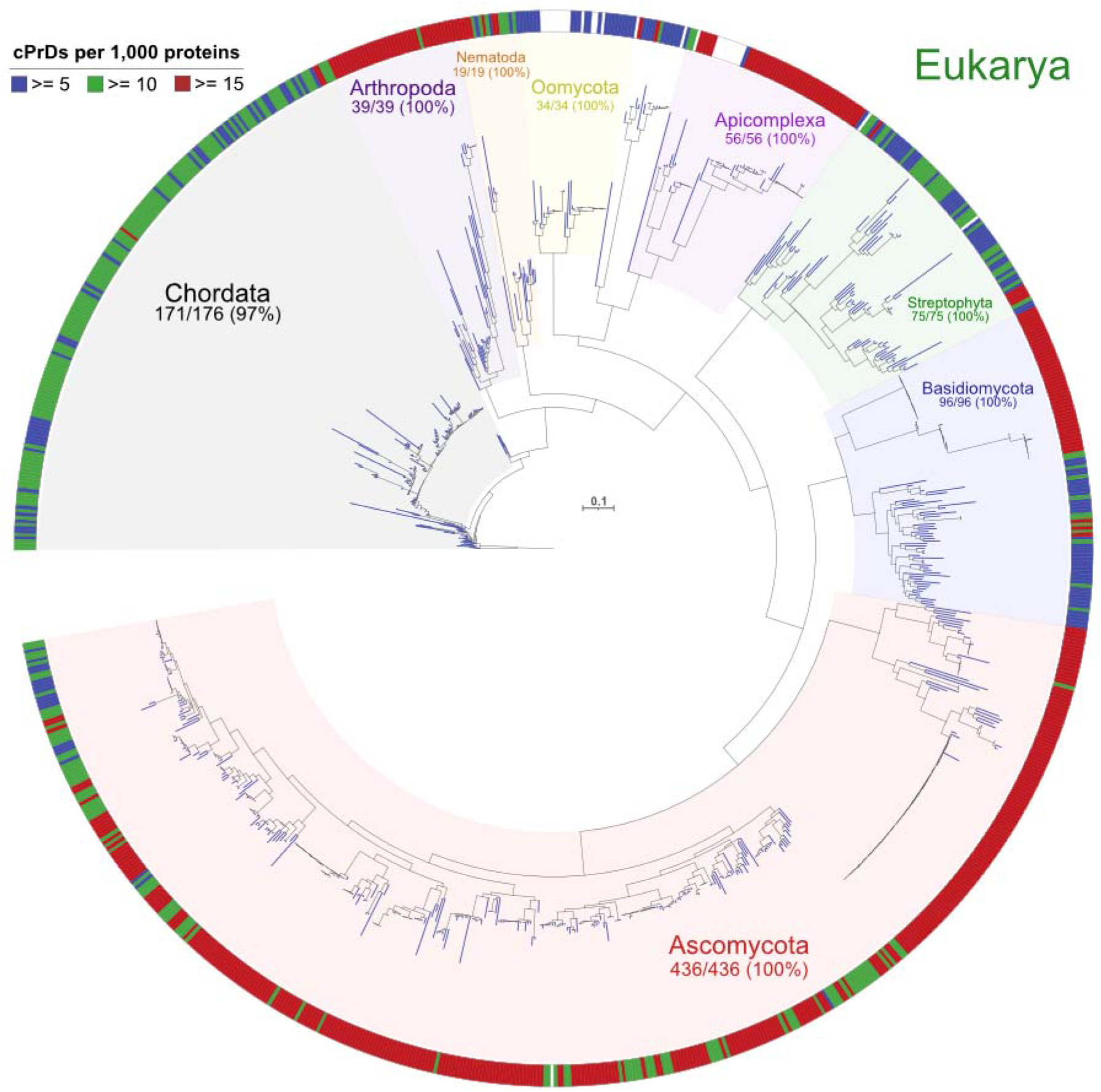
Phylogenomic tree of analyzed eukaryal proteomes with their distribution of cPrDs overlain. Tips with at least 1 cPRD identified are colored blue. The outer ring depicts those with greater than 5, 10, or 15 cPrDs per 1,000 proteins within a proteome.

### 3.2 Analysis of functions of prion candidates

To facilitate the identification of evolutionarily conserved prion functions within our dataset we leveraged the Gene Ontology (GO) annotations associated with all proteins in the UniProt database. GO assigns information to a protein in the context of 3 “namespaces” – molecular function (MF), biological process (BP), and cellular component (CC) – through a highly curated process involving both manual and automated methods (Ashburner et al. 2000). In addition to namespaces, GO terms are stored in a hierarchical structure of parent-child relationships and denoted as specific “depth” levels. Within a given namespace, a lower depth level (as in a lower number, for example, a depth of 1) will typically denote a broader, less specific annotation than a GO term with a depth level of 2. GO annotations can be thought of more as protein-domain level functional annotations than as full-protein functional annotations. To complement the GO annotations in our analysis of functions associated with cPrDs, we also annotated cPrDs with Kyoto Encyclopedia of Genes and Genomes (KEGG) (Kanehisa and Goto 2000) KEGG Orthology (KO) terms – which are more akin to full-protein functional annotations.

#### 3.2.1 Enriched GO terms in prion candidates

Within each domain of life, we tested for enrichment or purification (see Methods) of specific GO terms in our identified cPrD-containing proteins as compared with all of the proteins scanned. In the archaeal domain, 51 GO terms were found to be significantly enriched (meaning these specific GO terms were more likely to be found in a protein with an identified candidate prion-domain than in a protein without one), and 345 significantly purified (meaning these specific GO terms were more likely to be associated with a protein that did not have an identified cPrD; based on a Benjamini-Hochberg false-discovery rate of ≤ 0.05; Table S3). For Bacteria, 248 were found to be significantly enriched, and 1,243 were significantly purified (Table S4). And in the Eukarya, 2,601 GO terms were found to be significantly enriched, and 6,413 significantly purified (Table S5).

First, we focused on enriched GO terms (those more likely to be found in proteins with cPrDs than in proteins without cPrDs) in order to pursue functions that are consistently associated with prions. Within the GO molecular function namespace, there were two GO terms that were found enriched in all 3 domains: helicase activity (GO:0004386) and calcium ion binding (GO:0005509). For the biological process and cellular component namespaces, more broad-level terms were the only ones enriched: macromolecule catabolic process (GO:0009057) and the root term of cellular component (GO:0005575).

There were 10 molecular function GO terms that were found enriched in both Bacteria and Archaea (without eukaryotes) (Table 1). Within the GO namespace biological process, biological adhesion, and metabolic process were found enriched. Protein ubiquitination (GO:0016567) and xylan catabolic process (GO:0045493) were the most specific descriptions found (depth 9). Within the cellular component namespaces in both Bacteria and Archaea we found integral components of the membrane (GO:0016021) and extracellular region (GO:0005576) to be enriched (Table S6).

**Table 1.** GO terms that are statistically more likely to be found with cPrDs than without in multiple domains.

| Domains with shared enriched GO terms | Shared enriched molecular function GO terms |
| --- | --- |
| Archaea, Bacteria, and Eukarya | helicase activity (GO:0004386)<br>calcium ion binding (GO:0005509) |
| Archaea and Bacteria | cysteine-type endopeptidase activity (GO:0004197)<br>serine-type endopeptidase activity (GO:0004252) |
|  | helicase activity (GO:0004386)<br>hydrolase activity, hydrolyzing O-glycosyl compounds (GO:0004553)<br>calcium ion binding (GO:0005509)<br>ATP-dependent activity, acting on DNA (GO:0008094)<br>hydrolase activity, acting on glycosyl bonds (GO:0016798)<br>carbohydrate-binding (GO:0030246)<br>endo-1,4-beta-xylanase activity (GO:0031176)<br>xylanase activity (GO:0097599) |

**Table 2.** Numbers of candidates as a function of PLAAC COREScore for enriched GO Terms common to candidate prions in all three domains of life. The higher the COREScore, the higher the predicted prion-like behavior, making them likely better targets for experimental verification. Table S9 holds PLAAC results and sequences for just these candidates.

| GO Term | Domain | COREScore < 42 | COREScore >= 42 |
| --- | --- | --- | --- |
| GO:0005509<br>calcium ion binding | Archaea | 119 | 37 |
|  | Bacteria | 115 | 2 |
|  | Eukarya | 2,161 | 82 |
| GO:0004386<br>helicase activity | Archaea | 38 | 3 |
|  | Bacteria | 166 | 40 |
|  | Eukarya | 2,758 | 56 |

#### 3.2.2 Shared GO terms in prion candidates across domains

Among GO annotations that overlap all three domains of life, not focusing on enriched or not, we identified 84 that were common to all three domains (54 molecular functions, 22 biological process, 8 cell components) (Figure 4).

**Figure 4.**
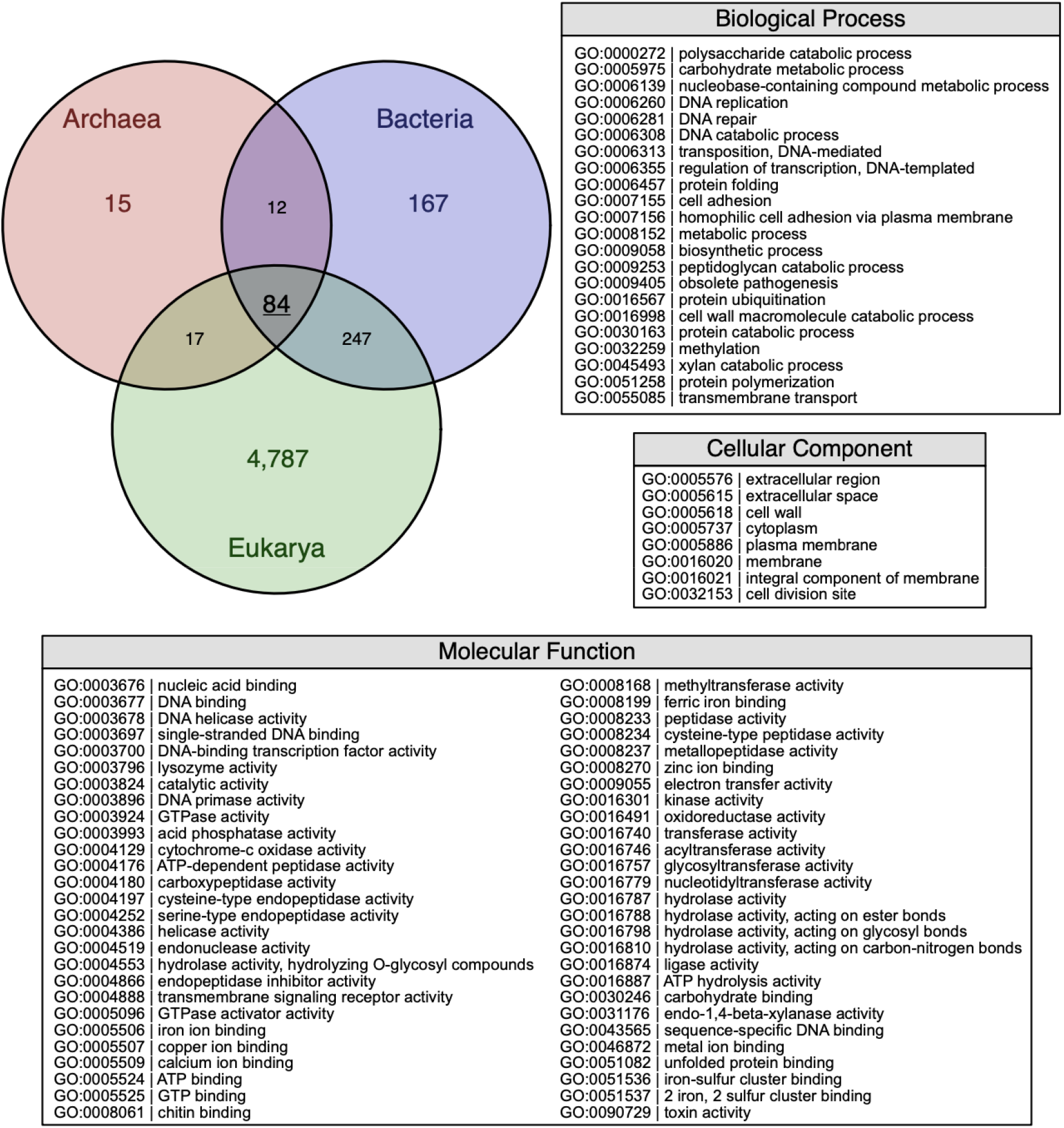
Shared GO terms of prion candidates across domains. The Venn diagram displays counts of GO terms in the same protein with candidate prion domains, including counts shared between domains (Table S7). The text depicts the 84 that are shared between all 3 domains in their given GO namespaces.

In all GO namespaces (molecular function, biological process, cellular component) we found a relatively low number of candidate prion proteins detected in archaea (e.g., Figure 1). This could be explained by a low number of archaea proteomes available in the Uniprot database, presumably due to difficulty in establishing laboratory conditions for culturing.

Within the GO molecular function namespace overlapping all three domains we found 82,489 candidate prion proteins: 533 in archaea, 3,667 in bacteria, and 78,289 in eukaryotes (Table S7). There were 54 different overlapping GO molecular function annotations. Among them, the most common annotations were: serine-type endopeptidases (GO:0004252) 15 archeal, 273 bacterial, 244 eukaryotic; transcription factors (GO:0003700) 2 archeal, 8 bacterial, 7,356 eukaryotic; single-stranded DNA binding proteins (GO:0003697) 1 archeal, 191 bacterial, 687 eukaryotic; kinases (GO:0016301) 1 archaeal, 49 bacterial, 569 eukaryotic.

Within the GO biological process namespace overlapping all three domains we found 10,073 candidate prion proteins: 254 in archaea, 1209 in bacteria, and 8,610 in eukaryotes (Table S7). There were 22 different overlapping GO biological process annotations. Among them, the most common annotations were: protein ubiquitination (GO:0016567) 133 archaeal, 6 bacterial, 176 eukaryotic (many of the archaeal/bacterial proteins associated with this GO term were annotated with K07218, nitrous oxidase accessory protein; see Table S7); cell adhesion (GO:0007155) 38 archaeal, 259 bacterial, 180 eukaryotic; DNA replication (GO:0006260) 1 archaeal, 225 bacterial, 82 eukaryotic; DNA-templated regulation of transcription (GO:0006355) 6 archaeal, 18 bacterial, 5351 eukaryotic; DNA repair (GO:0006281) 6 archaeal, 127 bacterial, 1345 eukaryotic.

Within the GO cellular component namespace overlapping all three domains we found 34,084 candidate prion proteins: 924 in archaea, 6,615 in bacteria, and 26,545 in eukaryotes (Table S7). There were 22 different overlapping GO cellular component annotations. Among them, the most common annotations were: integral component of membrane (GO:0016021) 842 archaeal, 5059 bacterial, 13569 eukaryotic; plasma membrane (GO:0005886) 3 archaeal, 406 bacterial, 1915 eukaryotic; cell wall (GO:0005618) 3 archaeal, 49 bacterial, 170 eukaryotic.

## 4. Discussion

### 4.1 Overlap of enriched GO terms of prion candidates across all three domains of life

We hypothesized that some functions associated with prions are conserved, along with prion domains, across different domains of life – allowing for the possibility that prion domains are essential to these specific GO-term functions. To pursue this hypothesis, we first focused on enriched GO term functions that were shared by candidate prion proteins (cPrPs) across all three domains of life. Finding those that span all three domains gives stronger credence to the possibility that these functions are truly conserved alongside prions rather than being a product of convergent evolution – though, importantly, convergent evolution is certainly not eliminated as a possibility. To expand the search beyond just what is shared among all three domains, we also focused on what is shared just between Bacteria and Archaea, as these targets may still give us insight into distantly related functions associated with prion domains.

#### 4.1.1 Overlap of enriched GO terms assigned to prion candidates across three domains of life

We identified two molecular functions GO terms that were found enriched in all three domains: helicase activity (GO:0004386) and calcium ion binding (GO:0005509; Figure 5). To assess how likely a prion candidate is to behave as a prion experimentally, PLAAC assigns a COREScore value. We used a COREScore value equal to 42 as an indication of a strong candidate because bacterial protein Rho acquired this value and was confirmed to form a prion in experiments (Yuan and Hochschild 2017; Pallarès and Ventura 2017). We think these are strong candidates to pursue experimentally. Table S9 holds PLAAC results and sequences on just these candidates.

**Figure 5.**
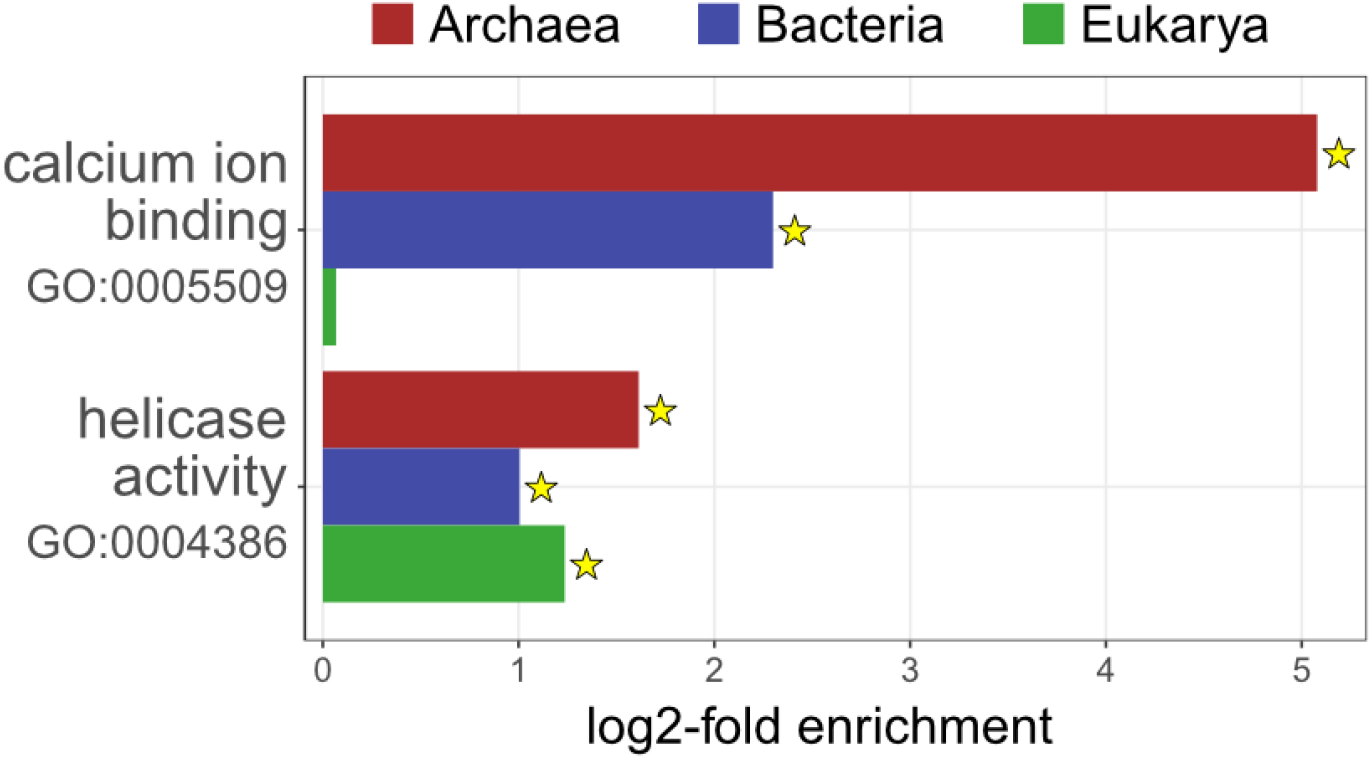
Log2-fold enrichment values for the 2 GO terms found to be enriched in cPrPs in all 3 domains. Those with stars had Benjamini-Hochberg adjusted-p-values ≤ 4E-8; for Eukarya, GO:0005509 had a BH adjusted p-value of 0.073.

We also looked at the KO annotations associated with candidate prion proteins (cPrPs) that held enriched GO annotations. KO annotations are more akin to whole protein-level annotations; while GO annotations are more akin to protein-domain level annotations. The group of prion candidates annotated with the calcium ion binding GO term (GO:0005509) was annotated with a variety of different KO functions across Archaea, Bacteria, and Eukarya, but the one consistent KO function that occurred in all three domains was K13974, a calcium-binding protein (Table S10; full KO annotation information for all cPrPs is available in our figshare repository linked in Methods).

Among candidates annotated with helicase activity (GO:0004386), the majority of bacterial prion candidates were annotated as K03628, transcription termination factor Rho. Transcription termination factor Rho was originally identified as a candidate prion protein by Iglesias et al. (Iglesias, de Groot, and Ventura 2015) and was later shown to form a self-perpetuating prion state in *Escherichia coli* (Yuan and Hochschild 2017). For Archaea, the most common KO annotation within this group of proteins was K11927, ATP-dependent RNA helicase RhlE – which was also found among bacterial prion candidates. In Eukarya, multiple different helicases showed up as prion candidates including ATP-dependent RNA helicase DDX6/DHH1 (K12614), DDX5/DBP2 (K12823), and DDX3X (K11594); Table S10). All these helicases are good candidates for experimental verification of prion properties. This observation is further strengthened by the fact that DDX5 was recently shown to form cytoplasmic aggregates in the brains of old killifish and mice. When studied in a yeast-based heterologous system, DDX5’s prion-like domain allowed these aggregates to propagate across many generations (Harel et al. 2022).

Prion proteins are known to be involved in stress response, and helicases are particularly good candidates for actuators of prion-based response as they can integrate diverse inputs and activate diverse outputs. Helicases provide immediate access to genetically complex traits by influencing the activity of many genes at the same time - in consequence enabling phenotypic variation that might be necessary for survival in a changing environment. The fact that helicases are one of only two groups of enriched functions of putative prions shared by all three domains of life supports the idea that prion mechanisms may be essential to population-level survival and hence might be evolutionarily conserved.

#### 4.1.2 Overlap of enriched GO terms of prion candidates between archaea and bacteria

As mentioned above, focusing on cPrD-associated functions that span all three domains helps support the notion that these are evolutionarily conserved across all of known life but it also limits our scope. To expand our search, we next focused on what is shared just between Archaea and Bacteria, as these targets too may provide a window into distantly related, but conserved, functions associated with prion domains (see Table 3). Table S8 GO terms enriched in both Bacteria and Archaea. When we searched for common GO annotations in these groups we found an overlap of GO terms in the biological processes namespace involved in adhesion, metabolic process, and protein modification (see Figure 6). Overlapping GO terms in the molecular function’s namespace of prion candidates were in agreement with these above-listed GO biological process terms. For example, the carbohydrate-binding function corresponded to the adhesion process. Enzymatic functions corresponded to the metabolic process, and protein ubiquitination corresponded to protein modification. And, of course, as enriched GO terms for helicase activity and calcium ion binding were shared between all 3 domains, those are also shared between Bacteria and Archaea. According to our dataset, based on this approach, these processes and functions are the most conserved prion functions in nature.

**Table 3.**
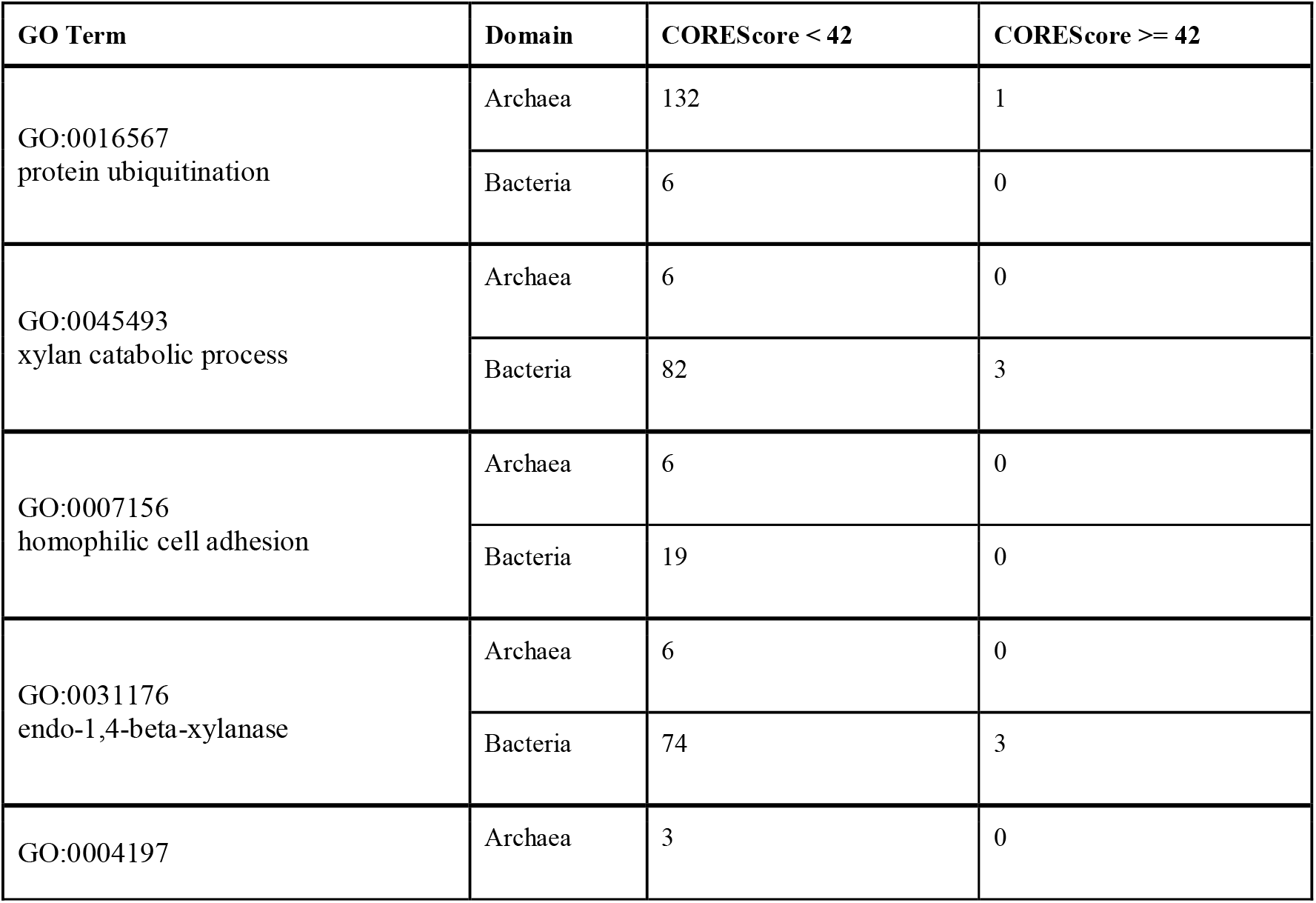
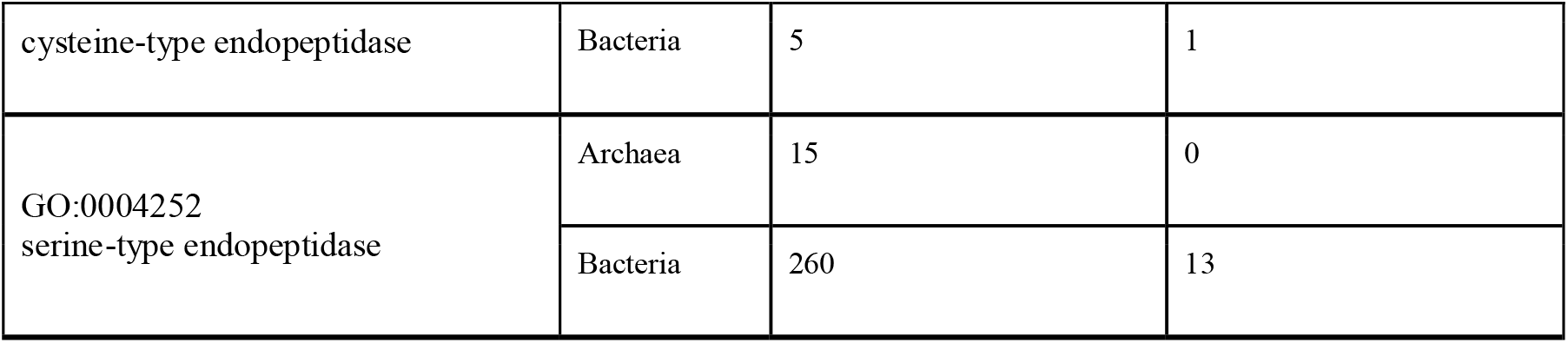
Numbers of candidates as a function of PLAAC COREScore for selected enriched GO Terms common to Archaea and Bacteria. The higher the COREScore, the higher the predicted prion-like behavior, making them likely better targets for experimental verification. Table S11 holds PLAAC results and sequences for just these candidates.

**Figure 6.**
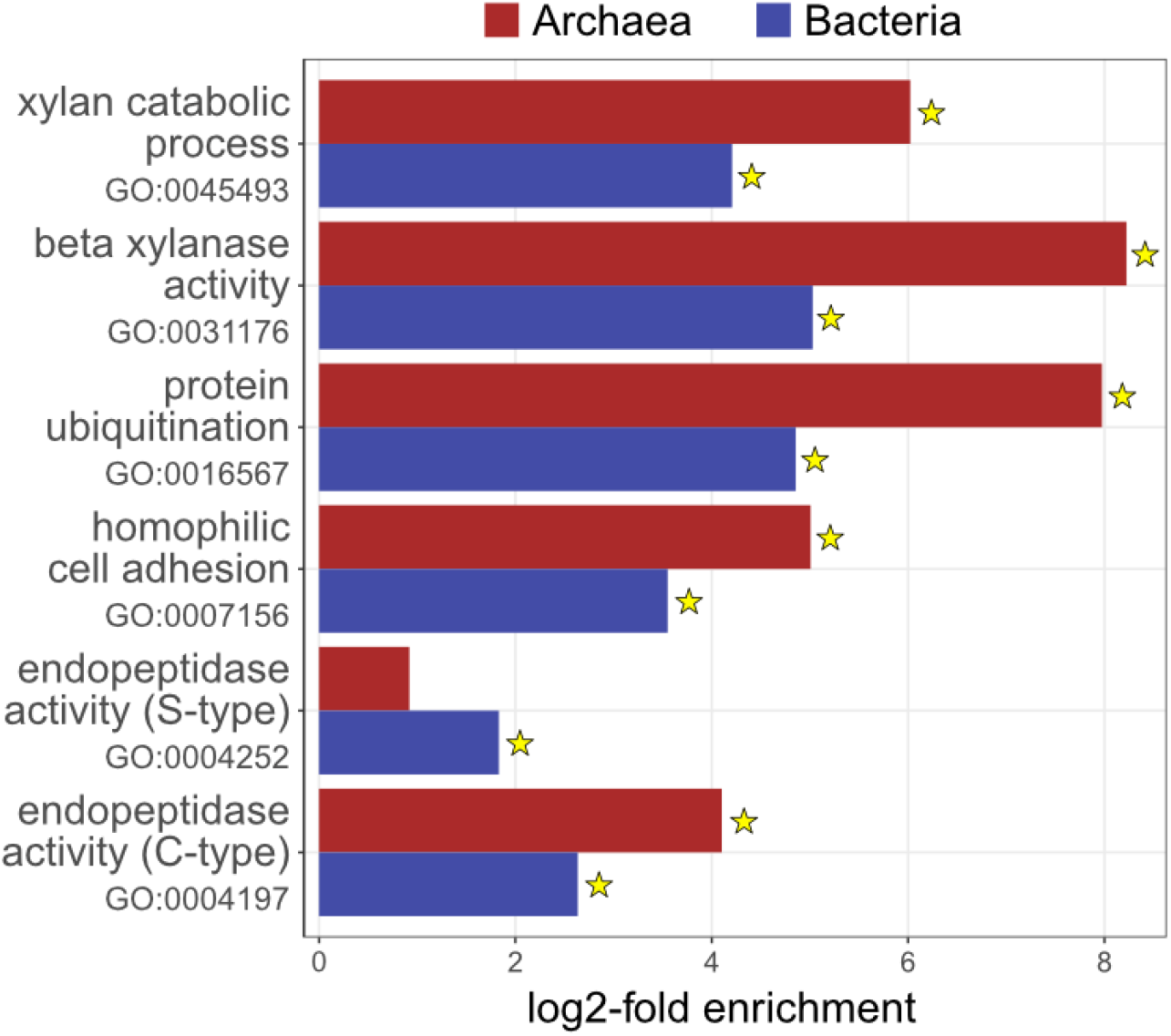
Log2-fold enrichment values for the 2 GO terms found to be enriched in cPrPs in both Bacteria and Archaea domains as compared to all proteins from those domains. Those with stars had Benjamini-Hochberg adjusted-p-values ≤ 0.012; for Archaea, GO:0004252 had a BH adjusted p-value of 0.19. “S-type” refers to serine-type; “C-type” refers to cysteine-type.

#### 4.1.3 Organisms harboring enriched GO terms that overlap across all three domains of life

After identifying enriched GO terms that overlap across all three domains of life, we wanted to see if there were any trends in the distribution of candidate prions associated with these specific GO terms with regard to the phylogenetic relationships of the organisms containing them. Presumably, it is possible that a cPrD being consistently associated with a specific GO term is either a consequence of that organism’s evolutionary history (which would present largely as monophyletic phylogenetic clades of organisms possessing the protein holding cPrD and GO term), horizontal gene transfer, or it could be due to independently evolved, yet common, characteristics (which, as with horizontal gene transfer, might present as a highly polyphyletic distribution).

For both the GO annotations calcium ion binding (GO:0005509) and helicase activity (GO:0004386) in Archaea, most candidates were detected within one monophyletic clade of Euryarchaeota, with a few others spread around members of the DPANN group and within the Thaumarchaeota (Figures S1 and S2). Considering the possibility of the ancient origin of prions one might expect to discover candidate prion proteins (cPrPs) annotated with these two GO terms within the superphylum Asgard -suspected progenitors of Eukaryota (Zaremba-Niedzwiedzka et al. 2017). The lack of candidate prions annotated with enriched GO terms overlapping all three domains (calcium ion binding and helicases) in Asgard archaea might be explained by the fact that this group consists mostly of uncultured and relatively understudied organisms, which lowers their representation in standard databases. Indeed, the UniProt reference proteome database we were working with here only included 16 from the Asgard group, all belonging to the candidate phylum Lokiarchaeota (Table S1). There is also the potential that the retained protein similarity of potentially homologous functions was too divergent to be functionally annotated with the same GO term.

Distribution patterns in Bacteria and Eukarya were similar for both of the conserved, enriched GO terms (calcium ion binding (GO:0005509) and helicase activity (GO:0004386)). In bacteria, we found prion candidates with associated calcium ion binding annotations in all major phyla, with a higher concentration of calcium ion binding within the Alphaproteobacteria, and a higher concentration of helicase activity within Actinobacteria (Figures S1 and S2). And in Eukarya, candidates with both GO terms are found roughly throughout the entire tree, with the exception being within the Chordata (Figures S1 and S2).

### 4.2 Overlap of GO terms of prion candidates across all three domains of life

Next, we identified GO annotations of prion candidates common to different domains of life whether they were statistically enriched in prion candidates over all proteins or not (meaning, now no longer focusing on “enriched” as was done above). Having an enriched molecular function, biological process, or cellular component (the GO namespaces) implies that a prion candidate is more likely to be associated with a certain GO term as compared to a protein that is not a prion candidate. That view helps focus on GO terms that seem to be only found in prion candidates, but it would miss those that can be with or without a candidate prion domain. But even without looking at enrichment, the proteins identified by PLAAC are valid prion candidates and, if their GO annotations are found to be common across all three domains of life, they may still represent some of the oldest prions on Earth. In this section, we focused on such prion candidates whose GO terms were not necessarily enriched, but were detected in all three domains of life.

#### 4.2.1 Overlap of molecular function GO terms of prion candidates across all three domains of life

Among GO molecular functions of candidate prion proteins that overlap across all three domains of life, the best represented in our dataset were transcription factors, kinases, DNA binding, peptidases, ribonucleotide binding, and metal binding (Figure 4; Table S7).

One prime example of this is DNA-binding transcription factor activity (GO:0003700). Transcription regulation is commonly implicated with prion biology of confirmed prion proteins, such as Ure 2 - [*URE3*^+^] prion (Wickner 1994), Mot3 - [*MOT3*^+^] prion (Alberti et al. 2009), and Sfp1 - [*ISP*^+^] prion (Volkov et al. 2002; Rogoza et al. 2010). Binding to nucleic acids is one of the most fundamental activities of life and plays a role in each step of the central dogma of biology. When we looked at KO annotations of prion candidates that fall under GO:0003700, we found that archaea and eukaryotes share the KO term, K21042, “HCMV protein UL11” (Table S12), which is also present in viruses. The candidate among archaea belongs to the genus *Thermoproteus*, a thermophile, whose protein contains the marR-type HTH domain, which is responsible for antibiotic resistance. Production of this protein may be a response to the presence of organisms in the environment that produce antibiotic-like substances (e.g., fungi). The same protein of the UL11 type in eukaryotes has the FHA domain that takes part in an ancient and widespread mechanism of regulation based on the phospho-dependent assembly of protein complexes (Durocher and Jackson 2002). In bacteria, prion candidates annotated as DNA-binding transcription factors (GO:0003700) consisted mostly of RNA polymerase sigma-70 factor, ECF subfamily (K12888) which is the major factor that initiates transcription in bacteria.

Single-stranded DNA binding (GO:0003697) has been identified as common among prions not only in our dataset but also by Harrison and Paul (P. M. Harrison 2019). Single-strand binding proteins are exceptionally important for maintaining the stability of an organism’s genome, as they are involved in key processes taking place in the nucleus, such as DNA replication, repair, and recombination. Among single-stranded DNA binding candidate prion proteins, we found 190 annotated as K03111, single-strand DNA-binding protein, but only in bacteria. In eukaryotes we identified 3 other KO descriptions: K21390, adhesion defective protein 2, which is responsible for the transcriptional regulation of cell adhesion; K12888, heterogeneous nuclear ribonucleoprotein U, intra-nuclear proteins that take part in many processes of the nucleus metabolic pathway including organization of chromatin, regulation of telomere length, transcription, and alternative mRNA splicing; and K13184, ATP-dependent RNA helicase A, a multifunctional protein involved in processes such as DNA replication, post-transcriptional regulation of RNA, mRNA translation, and silencing. All KO term groups identified single-stranded DNA binding proteins play a particularly important role in keeping DNA and RNA functioning properly. As for the archaea, the only prion candidate from this GO did not have any KO term assigned.

Kinase activity (GO:0016301) was another GO annotation that overlapped all three domains. Within this GO annotation, we found diverse kinase-related proteins based on their KO annotations including activators of two-component systems (e.g. K20340), serine/threonine kinase activators (e.g., K08286), and activators of tyrosine kinases (eK23453) – with annotations virtually only being ascribed to eukarya (Table S12). These have a diverse repertoire of functions ranging from regulating cell death, cell migration, and cell adhesion, to general transcriptional repression related to circadian rhythm.

Another group of candidate prion proteins that overlapped all three domains were serine-type endopeptidases (GO:0004252). Peptidases are active enzymes that cleave peptide bonds in proteins and peptides by hydrolysis, and serine-type endopeptidases fall into a class of peptidases that are characterized by the presence of a serine residue in the active site of the enzyme. They are of extremely widespread occurrence from prokaryotes to vertebrates and exhibit diverse functions (Kraut 2003). When we looked at KO annotations of prion candidates that fall under the serine-type endopeptidases (GO:0004252), in bacteria we found many annotated as K08372 - putative serine protease PepD (Table S12). Most PepD peptidases have PDZ domains that recognize and process misfolded proteins at the cell membrane, leading to the activation of signaling pathways and the establishment of a feedback loop that can facilitate bacterial adaptation (White et al. 2010; Muley et al. 2019). This observation is consistent with the hypothesis that prion formation might be an ancient mechanism facilitating adaptation. Among endopeptidases in archaea, we found ATP-dependent Lon protease. This protease is found both in mitochondria and bacteria and shares high similarities between the organelle and the domain (Wang et al. 1993). Among eukaryotic prion candidates annotated with the GO term serine-type endopeptidases (GO:0004252), we found rhomboid proteases (e.g., K19225; Table S12) that are common in all domains of life and are implicated in various functions including cell signaling, quorum sensing, and homeostasis (Urban and Dickey 2011). Recently, Rhomboid Protease RHBDL4 was shown to cleave amyloid precursor protein (APP) a key molecule in the etiology Alzheimer’s disease (Paschkowsky et al. 2016; Penalva et al. 2021; Urban and Moin 2014).

Another GO description overlapping all three domains of life was ribonucleotide binding (GO:0032553), which similarly to the DNA-binding transcription factor activity (GO:0003700), is central to biology because ribonucleotides are primary sources of energy for biochemical reactions. Ribonucleotide binding is a parent term of four GO annotations that also was associated with candidate prions in our dataset in all three domains: ATP binding (GO:0005524), ATP hydrolysis activity (GO:0016887), GTP binding (GO:0005525), and GTPase activity (GO:0003924).

The last major group of annotations common to all domains (regardless of enrichment) identified in our dataset was annotated as metal-binding proteins (GO:0046872), which play essential roles in a wide range of structural and catalytic functions. Similar to the above-mentioned functions these are also central to all biology. Child terms of GO:0046872 that are also common to all domains were zinc ion binding (GO:0008270; 1 archeal, 94 bacterial, 10781 eukaryotic), iron ion binding (GO:0005506; 1 archeal, 9 bacterial, 38 eukaryotic), copper ion binding (GO:0005507; 25 archeal, 17 bacterial, 88 eukaryotic), and ferric iron-binding (GO:0008199; 1 archeal, 1 bacterial, 1 eukaryotic). KO annotations for the candidate prion proteins holding these shared functions were highly varied (Table S12).

#### 4.2.2 Overlap of biological process GO terms of prion candidates across all three domains of life

When we analyzed GO terms within the biological process namespace that overlapped across three domains of life, we found that for archaea, the greatest number of prion candidates were annotated with protein ubiquitination (GO:0016567; 133 proteins; Table S7). For bacteria, two GO terms were more common than others: cell adhesion (GO:0007155; 259 proteins) and DNA replication (GO:0006260; 225 proteins). For eukaryotes, the greatest number of prion candidates were annotated as DNA-templated regulation of transcription (GO:0006355; 5,351 proteins; Table S7).

Another GO annotation brought to our attention was DNA repair (GO:0006281), which was previously noted as an abundant biological process among candidate prions by (Iglesias, de Groot, and Ventura 2015). This annotation was clustered by these authors in the group stimulus-response process with some candidates annotated specifically as a cellular response to DNA damage stimulus (GO:0006974).

Other GO terms that were detected in our analysis as well as others (Iglesias, de Groot, and Ventura 2015), were grouped under a common category of invasion and virulence (GO:0009405, obsolete pathogenesis, and GO:0000272 polysaccharides catabolic process), and under another broad category of nucleotide metabolism (GO:0006353, DNA-templated transcription termination and GO:0006260, DNA replication).

One of the major groups of prion candidates identified by (Iglesias, de Groot, and Ventura 2015) was annotated as GO:0009056, catabolic process. Many catabolic processes were also present in our dataset: GO:0000272, polysaccharide catabolic process; GO:0006308, DNA catabolic process; GO:0045493, xylan catabolic process; GO:0030163, protein catabolic process; GO:0000272, polysaccharide catabolic process; GO:0009253, peptidoglycan catabolic process; and GO:0016998, cell wall macromolecule catabolic process. The last two GO annotations (GO:0009253 and GO:0016998) have also been described as common annotations of bacterial prion candidates by (P. M. Harrison 2019).

A high number of candidates in archaea and bacteria candidates were annotated as GO:0007155 cell adhesion (38 archaeal, 259 bacterial), which might be related to the fact that both groups of organisms often form biofilms.

#### 4.2.3 Overlap of cellular component GO terms of prion candidates across all three domains of life

In our dataset the majority of candidate prion proteins were annotated as an integral component of membranes (GO:0016021). Localization of multiple candidate prion proteins on the periphery of the cell was noticed by other authors as well (P. M. Harrison 2019; Batlle et al. 2017). In addition to those we found a surprisingly high number of candidate prion proteins annotated as proteins occupying extracellular region - not attached to the cell surface (GO:0005576). So far no extracellular protein was found to form prions but taking into account a high number of candidate prion proteins annotated in this region it is likely that some of them could become the first extracellular prions verified experimentally.

### 4.3 Could ancient prions facilitate adaptation?

Prions have been hypothesized to facilitate generating diverse responses to changes in environmental or cellular conditions by regulating gene expression patterns (Shorter and Lindquist 2005). In yeast, prion [*GAR*^+^] regulates whether cells are metabolic specialists or generalists (Brown and Lindquist 2009), prion [*SMAUG*^+^] regulates sporulation (Chakravarty et al. 2020), and prion [*ESI*^+^] influences the activity of multiple genes by regulating the expression of subtelomeric regions in response to environmental stresses such as antifungal drugs (Harvey et al. 2020). In Bacteria, prion-aggregation of protein Rho influences the expression of hundreds of genes, dramatically influencing cell phenotypes, potentially rendering prion-harboring cells better suited to rapid changes in the environment (Yuan and Hochschild 2017). Some of the functions of candidate prion proteins identified in our dataset that overlap across all three domains of life (namely transcription factors, DNA binding, and peptidases) are particularly well suited to generating diverse responses to changes in their environment. We suspect that these potential prions can provide a rapid means for organisms to explore the evolutionary landscape, and, given their presence across all three domains, they may be some of the most ancient functions associated with prions.

Perhaps one of the most efficient ways of generating diverse responses to changes in environmental or cellular conditions is by changing signal response and regulation pathways. Candidate prions with associated kinase-related functions are therefore intriguing to be found across all three domains of life in our dataset (Figure 4; GO:0016301). Generally, the potential role of candidate prions in signal transduction and regulation suggests they may have played a critical role in the evolution of some of the complex pathways that have allowed life to expand into a wide range of environmental niches.

Indeed, a question that has plagued the study of kinases is how such complexity could have evolved and how orthologous signaling proteins diverge (Capra and Laub 2012). Suspending the pressures of natural selection acting on a protein for several generations following a duplication event would allow for a sufficient number of step-wise mutations to accumulate, leading to the large mutational jumps required for preventing cross talk between newly diverged kinases and their ancestors. Because prion evolution can in part be dictated by both DNA and protein heritability (Wickner and Kelly 2016), they may also provide a partial solution to the question of how newly evolved signaling proteins prevent cross-talk with their very similar ancestors. If a duplicated signaling protein becomes functionally inert due to being aggregated into fibrils in a prionic form, a larger number of mutations are allowed to accumulate, independent of selection. Because these prionic forms can be inherited by daughter cells via the phenomenon of seeding, this state of non-selective mutagenesis can carry on in daughter cells for many generations before the aggregated prion state ends and function is restored. This mechanism allows for large mutational “jumps” that could lead to large enough divergence to prevent cross-talk with orthologous ancestral signaling proteins – being one example of how proteins could accumulate many step-wise mutations without selection acting on individual mutations along the way (See Figure 5). Mechanisms for accelerated evolution of kinase proteins have been theorized as ways of preventing cross-talk, such as diversity generating retroelements, specific to the cyanobacterial phylum (Vallota-Eastman et al. 2020). However, these retroelements have only been found in viruses and prokaryotes where they usually target genes encoding for proteins involved in cell-cell or phage-cell attachment. Cyanobacteria are the only phylum where they have been found to target kinase genes specifically.. Prionic kinases, being found in all domains of life, are thus an intriguing potential mechanism for circumventing stepwise mutation. Furthermore, the kind of conformational switching that the prion state provides proteins allows for multiple activity states from the same polypeptide sequence, providing a greater efficiency and economy for the proteome.

Signal response is evolutionarily ancient as it is essential for multicellularity and would even be needed in LUCA to mount a simple metabolic response to changes in nutrients or increases in environmental toxicity (e.g., metal concentration, pH, etc.). While the role of prions in allowing for the greater divergence of signaling proteins remains to be explored, it is clear that they are important for this evolutionarily ancient mechanism of species adaptation.

## 5. Conclusions

In proteomes of organisms representing three different domains of life, we identified candidate prion domains and then analyzed the functional annotations associated with the proteins harboring them. Helicases and calcium ion binding proteins were enriched among prion protein candidates in all three domains of life – meaning these specific GO terms were more likely to be associated with a protein with an identified candidate prion domain than in a protein without one. Beyond just those enriched in candidate prions, we identified numerous functional annotations that are associated with candidate prions in all three domains. Some of the most represented included peptidases, transcription factors, single-stranded DNA binding proteins, and kinases; functions that are fundamental to the proper functioning of cells.

The role of prions in responses to changes in environmental conditions has been described as a process called biological “bet hedging” that can facilitate the survival of the resultant phenotypically heterogeneous population (Jong et al. 2011; Halfmann et al. 2010). In this model, the accumulation of prionic forms of proteins provides some phenotypic change that allows the cell, and in turn, some subset of the population, to potentially be better suited to a new environment. Aggregation of this prion form of the protein also typically renders the protein functionally inert with regard to its previous, more traditionally understood function, which could then allow for random genetic mutagenesis to occur more frequently, independent of selective pressures it otherwise would have faced. In these cases, due to chance, selection may then favor mutated progeny even once it is no longer experiencing prion aggregation (Figure 7). This is a mechanism that could allow for adaptation and population expansion into more diverse environments, and/or could allow for functional expansion of proteins involved in essential processes such as signal transduction and regulation.

**Figure 7.**
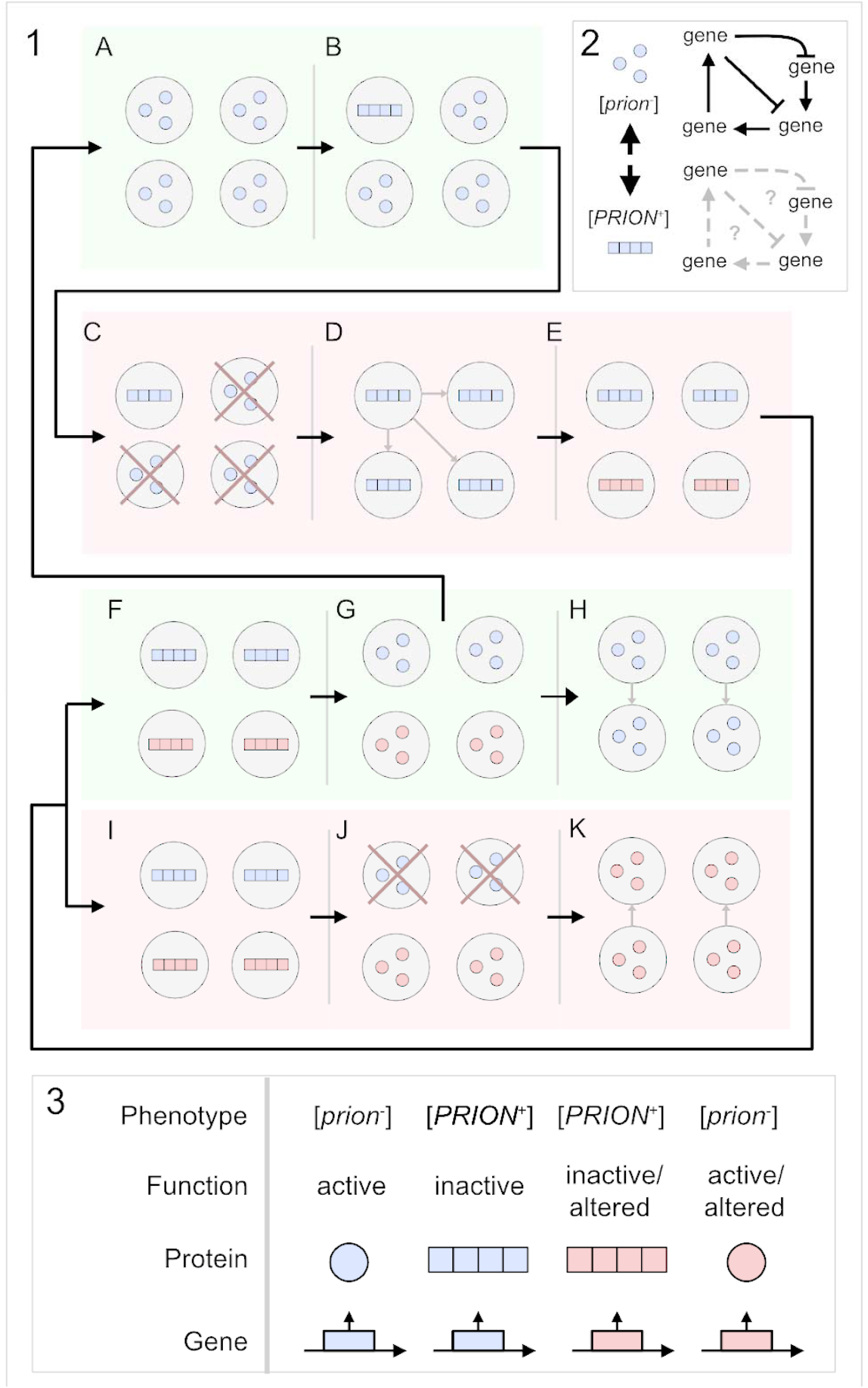
**1**. A. Population of cells in a given environment (green background). B. Spontaneous acquisition of prion phenotype. Squares represent amyloid fibril. C. Change of the environment (red background). Survival of the population is possible because it was phenotypically heterogeneous, the phenomenon is known as biological “bet hedging” - discussed in the text. In the new environment, the prion phenotype undergoes positive selection. D. Reversion to non-prion phenotype is counter-selected. Prion phenotype dominates the population. E. Because the protein product is locked in prion aggregate (inactive in this example) the gene coding for prion protein experiences less evolutionary pressure and more mutations are tolerated. As a result, the population diverges. There are now two genetically different but phenotypically identical populations of cells. The diversity is at the level of genes but for simplification, we marked the aggregates in two different colors. F. Environment reverses to the original state. Prion phenotype no longer experiences environmental selection and with time will be reduced. G. Prion is lost due to natural selection and phenotype reverses to one that depends on the active form of the prion protein. The mutations acquired in point F can either experience negative selection pressure or, if benevolent, will help adapt to the new environment when encountered again in the future. H. The original population is restored back to the state from point A. I. An alternative scenario in which the environment does not reverse to the original state and selection pressure on prion phenotype continues. J. Reversion to non-prion phenotype is still counter-selected but the extended time during which new mutations can accumulate eventually leads to the emergence of protein that facilitates survival in the new environment - even when not aggregated in the form of prion. K. Eventually the population survives even when the prion phenotype is lost. The population is now adapted to the new environment. 2. Prion conversion influences regulatory gene cascades influencing the expression of many genes at once, providing a rapid means for organisms to explore the evolutionary landscape. 3. Relationship between gene, protein, function, and phenotype expained with symbols used in panel 1.

The exciting possibility that these functions are also subjected to regulation through prion-formation remains to be verified experimentally. Many of the proteins identified in our study had high COREscore values, indicative of stronger predicted prion-like behavior, making them excellent targets to pursue experimentally. Confirming that these proteins can form prions would indicate that at least some essential protein functions are accompanied by prion domains across great evolutionary distances. If that turns out to be true, the role of prions in the regulation of the fundamental cell processes could be an evolutionarily ancient one, even if individual prion domains themselves are not evolutionarily conserved.

Based on our results, we hypothesize that prions that can influence the expression of many genes at once provide a rapid means for organisms to explore the evolutionary landscape – and this might be the most ancient function of prions conserved across all domains of life.

## Supporting information

Supplemental materials list

## Acknowledgments

We acknowledge Blue Marble Space Institute of Science (BMSIS) for organizational support. We thank doctor Daniel Jarosz for his insightful comments during manuscript preparation. We thank Polish Astrobiology Society and members of the Prion Task Force (www.astrobio.pl/prion), especially Magdalena Pilska-Piotrowska for help in developing the concept of domesticated amyloids and prions. S.S would like to acknowledge the SETI Forward 2020 from the SETI Institute. The authors would also like to thank the editors for the invitation to participate in this special issue.

## Author Contributions

Conceptualization, T.Z, M.D.L, L.J.R; methodology, T.Z, M.D.L, S.S; formal analysis, M.D.L, S.S, M.M; investigation, T.Z, M.D.L, S.S, A.V-E, M.K, M.M; data curation, T.Z, M.K, M.M; writing—original draft preparation, T.Z, M.D.L, S.S, A.V-E, M.K, M.M; writing, review and editing, T.Z, M.D.L, S.S, A.V-E, L.J.R; visualization, M.D.L, S.S, A.V-E,; project administration, T.Z, M.D.L, L.J.R. All authors have read and agreed to the published version of the manuscript

## Institutional Review Board Statement

Not applicable.

## Informed Consent Statement

Not applicable

## Data Availability Statement

The code and data for reproducing the computational analyses performed herein are available at: https://figshare.com/projects/Zajkowski_et_al_2022_3-domain_prion_data_and_code_repository/133155

## References

Afsar Minhas, Fayyaz Ul Amir, Eric D. Ross, and Asa Ben-Hur. 2017. “Amino Acid Composition Predicts Prion Activity.” PLoS Computational Biology 13 (4): e1005465.

Alberti, Simon, Randal Halfmann, Oliver King, Atul Kapila, and Susan Lindquist. 2009. “A Systematic Survey Identifies Prions and Illuminates Sequence Features of Prionogenic Proteins.” Cell 137 (1): 146–58.

“Anaconda.” n.d. Anaconda. Accessed February 21, 2022. https://www.anaconda.com/.

An, Lu, and Paul M. Harrison. 2016. “The Evolutionary Scope and Neurological Disease Linkage of Yeast-Prion-like Proteins in Humans.” Biology Direct 11 (July): 32.

Aramaki, Takuya, Romain Blanc-Mathieu, Hisashi Endo, Koichi Ohkubo, Minoru Kanehisa, Susumu Goto, and Hiroyuki Ogata. 2019. “KofamKOALA: KEGG Ortholog Assignment Based on Profile HMM and Adaptive Score Threshold.” Bioinformatics 36 (7): 2251–52.

Ashburner, Michael, Catherine A. Ball, Judith A. Blake, David Botstein, Heather Butler, J. Michael Cherry, Allan P. Davis, et al. 2000. “Gene Ontology: Tool for the Unification of Biology.” Nature Genetics 25 (1): 25–29.

Batlle, Cristina, Valentin Iglesias, Susanna Navarro, and Salvador Ventura. 2017. “Prion-like Proteins and Their Computational Identification in Proteomes.” Expert Review of Proteomics 14 (4): 335–50.

Beyersdörfer, Till. 2019. Curli Expression and Biofilm Formation by Escherichia Coli Isolates.

Bissig, Christin, Leila Rochin, and Guillaume van Niel. 2016. “PMEL Amyloid Fibril Formation: The Bright Steps of Pigmentation.” International Journal of Molecular Sciences 17 (9). https://doi.org/10.3390/ijms17091438.

Bourbon, Henri-Marc. 2008. “Comparative Genomics Supports a Deep Evolutionary Origin for the Large, Four-Module Transcriptional Mediator Complex.” Nucleic Acids Research 36 (12): 3993–4008.

Brown, Jessica C. S., and Susan Lindquist. 2009. “A Heritable Switch in Carbon Source Utilization Driven by an Unusual Yeast Prion.” Genes & Development 23 (19): 2320–32.

Buchan, J. Ross, Denise Muhlrad, and Roy Parker. 2008. “P Bodies Promote Stress Granule Assembly in Saccharomyces Cerevisiae.” The Journal of Cell Biology 183 (3): 441–55.

Capella-Gutiérrez, Salvador, José M. Silla-Martínez, and Toni Gabaldón. 2009. “trimAl: A Tool for Automated Alignment Trimming in Large-Scale Phylogenetic Analyses.” Bioinformatics 25 (15): 1972–73.

Capra, Emily J., and Michael T. Laub. 2012. “Evolution of Two-Component Signal Transduction Systems.” Annual Review of Microbiology. https://doi.org/10.1146/annurev-micro-092611-150039.

Cascarina, Sean M., and Eric D. Ross. 2020. “Natural and Pathogenic Protein Sequence Variation Affecting Prion-like Domains within and across Human Proteomes.” BMC Genomics 21 (1): 1–18.

Chakrabortee, Sohini, James S. Byers, Sandra Jones, David M. Garcia, Bhupinder Bhullar, Amelia Chang, Richard She, et al. 2016. “Intrinsically Disordered Proteins Drive Emergence and Inheritance of Biological Traits.” Cell 167 (2): 369–81.e12.

Chakrabortee, Sohini, Can Kayatekin, Greg A. Newby, Marc L. Mendillo, Alex Lancaster, and Susan Lindquist. 2016. “Luminidependens (LD) Is an Arabidopsis Protein with Prion Behavior.” Proceedings of the National Academy of Sciences of the United States of America 113 (21): 6065–70.

Chakravarty, Anupam K., Tina Smejkal, Alan K. Itakura, David M. Garcia, and Daniel F. Jarosz. 2020. “A Non-Amyloid Prion Particle That Activates a Heritable Gene Expression Program.” Molecular Cell 77 (2): 251–65.e9.

Cieśla, Małgorzata, Ewa Makała, Marta Płonka, Rafał Bazan, Kamil Gewartowski, Andrzej Dziembowski, and Magdalena Boguta. 2015. “Rbs1, a New Protein Implicated in RNA Polymerase III Biogenesis in Yeast Saccharomyces Cerevisiae.” Molecular and Cellular Biology 35 (7): 1169–81.

Dobson, Christopher M. 2003. “Protein Folding and Misfolding.” Nature. https://doi.org/10.1038/nature02261.

Durocher, Daniel, and Stephen P. Jackson. 2002. “The FHA Domain.” FEBS Letters 513 (1): 58–66.

Eddy, Sean R. 2011. “Accelerated Profile HMM Searches.” PLoS Computational Biology 7 (10): e1002195.

Edgar, Robert C. 2004. “MUSCLE: A Multiple Sequence Alignment Method with Reduced Time and Space Complexity.” BMC Bioinformatics 5 (August): 113.

Ehsani, Sepehr, Renzhu Tao, Cosmin L. Pocanschi, Hezhen Ren, Paul M. Harrison, and Gerold Schmitt-Ulms. 2011. “Evidence for Retrogene Origins of the Prion Gene Family.” PloS One 6 (10): e26800.

Espinosa Angarica, Vladimir, Alfonso Angulo, Arturo Giner, Guillermo Losilla, Salvador Ventura, and Javier Sancho. 2014. “PrionScan: An Online Database of Predicted Prion Domains in Complete Proteomes.” BMC Genomics 15 (February): 102.

Espinosa Angarica, Vladimir, Salvador Ventura, and Javier Sancho. 2013. “Discovering Putative Prion Sequences in Complete Proteomes Using Probabilistic Representations of Q/N-Rich Domains.” BMC Genomics 14 (May): 316.

Fleming, Eleanor, Andy H. Yuan, Danielle M. Heller, and Ann Hochschild. 2019. “A Bacteria-Based Genetic Assay Detects Prion Formation.” Proceedings of the National Academy of Sciences of the United States of America 116 (10): 4605–10.

Garai, Sampurna, Citu, Sneh L. Singla-Pareek, Sudhir K. Sopory, Charanpreet Kaur, and Gitanjali Yadav. 2021. “Complex Networks of Prion-Like Proteins Reveal Cross Talk Between Stress and Memory Pathways in Plants.” Frontiers in Plant Science 12 (July): 707286.

Garcia, Melissa C., Janis T. Lee, Caleen B. Ramsook, David Alsteens, Yves F. Dufrêne, and Peter N. Lipke. 2011. “A Role for Amyloid in Cell Aggregation and Biofilm Formation.” PloS One 6 (3): e17632.

Gazit, Ehud. 2007. “Self Assembly of Short Aromatic Peptides into Amyloid Fibrils and Related Nanostructures.” Prion 1 (1): 32–35.

Gene Ontology Consortium. 2021. “The Gene Ontology Resource: Enriching a GOld Mine.” Nucleic Acids Research 49 (D1): D325–34.

Gil-Garcia, Marcos, Valentín Iglesias, Irantzu Pallarès, and Salvador Ventura. 2021. “Prion-like Proteins: From Computational Approaches to Proteome-Wide Analysis.” FEBS Open Bio 11 (9): 2400–2417.

Goncharoff, Dustin K., Zhiqiang Du, and Liming Li. 2018. “A Brief Overview of the Swi1 Prion-[SWI+].” FEMS Yeast Research 18 (6). https://doi.org/10.1093/femsyr/foy061.

Halfmann, Randal, Simon Alberti, and Susan Lindquist. 2010. “Prions, Protein Homeostasis, and Phenotypic Diversity.” Trends in Cell Biology 20 (3): 125–33.

Halfmann, Randal, and Susan Lindquist. 2010. “Epigenetics in the Extreme: Prions and the Inheritance of Environmentally Acquired Traits.” Science 330 (6004): 629–32.

Harel, Itamar, Yiwen R. Chen, Inbal Ziv, Param Priya Singh, Paloma Navarro Negredo, Uri Goshtchevsky, Wei Wang, et al. n.d. “Identification of Protein Aggregates in the Aging Vertebrate Brain with Prion-like and Phase Separation Properties.” https://doi.org/10.1101/2022.02.26.482115.

Harrison, Alice Ford, and James Shorter. 2017. “RNA-Binding Proteins with Prion-like Domains in Health and Disease.” Biochemical Journal 474 (8): 1417–38.

Harrison, Luke B., Zhan Yu, Jason E. Stajich, Fred S. Dietrich, and Paul M. Harrison. 2007. “Evolution of Budding Yeast Prion-Determinant Sequences across Diverse Fungi.” Journal of Molecular Biology 368 (1): 273–82.

Harrison, Paul M. 2019. “Evolutionary Behaviour of Bacterial Prion-like Proteins.” PloS One 14 (3): e0213030.

Harrison, Paul M., and Mark Gerstein. 2003. “A Method to Assess Compositional Bias in Biological Sequences and Its Application to Prion-like Glutamine/asparagine-Rich Domains in Eukaryotic Proteomes.” Genome Biology 4 (6): R40.

Harvey, Zachary H., Anupam K. Chakravarty, Raymond A. Futia, and Daniel F. Jarosz. 2020. “A Prion Epigenetic Switch Establishes an Active Chromatin State.” Cell 180 (5): 928–40.e14.

Holmes, Daniel L., Alex K. Lancaster, Susan Lindquist, and Randal Halfmann. 2013. “Heritable Remodeling of Yeast Multicellularity by an Environmentally Responsive Prion.” Cell 153 (1): 153–65.

Hug, Laura A., Brett J. Baker, Karthik Anantharaman, Christopher T. Brown, Alexander J. Probst, Cindy J. Castelle, Cristina N. Butterfield, et al. 2016. “A New View of the Tree of Life.” Nature Microbiology 1 (April): 16048.

Iglesias, Valentin, Natalia S. de Groot, and Salvador Ventura. 2015. “Computational Analysis of Candidate Prion-like Proteins in Bacteria and Their Role.” Frontiers in Microbiology. https://doi.org/10.3389/fmicb.2015.01123.

Itakura, Alan K., Anupam K. Chakravarty, Christopher M. Jakobson, and Daniel F. Jarosz. 2020. “Widespread Prion-Based Control of Growth and Differentiation Strategies in Saccharomyces Cerevisiae.” Molecular Cell 77 (2): 266–78.e6.

Jahn, Thomas R., and Sheena E. Radford. 2005. “The Yin and Yang of Protein Folding.” FEBS Journal. https://doi.org/10.1111/j.1742-4658.2005.05021.x.

Jarosz, Daniel F., and Vikram Khurana. 2017. “Specification of Physiologic and Disease States by Distinct Proteins and Protein Conformations.” Cell 171 (5): 1001–14.

Jarvis, Suzi, and Anika Mostaert. 2012. The Functional Fold: Amyloid Structures in Nature. CRC Press.

Jong, Imke G. de, Imke G. de Jong, Patsy Haccou, and Oscar P. Kuipers. 2011. “Bet Hedging or Not? A Guide to Proper Classification of Microbial Survival Strategies.” BioEssays. https://doi.org/10.1002/bies.201000127.

Kanehisa, Minoru, and Susumu Goto. 2000. “KEGG: Kyoto Encyclopedia of Genes and Genomes.” Nucleic Acids Research 28 (1): 27–30.

Kanehisa, M., Y. Sato, M. Kawashima, M. Furumichi, and M. Tanabe. 2016. “KEGG as a Reference Resource for Gene and Protein Annotation.” Nucleic Acids Research 44 (D1). https://doi.org/10.1093/nar/gkv1070.

Kato, Masato, Tina W. Han, Shanhai Xie, Kevin Shi, Xinlin Du, Leeju C. Wu, Hamid Mirzaei, et al. 2012. “Cell-Free Formation of RNA Granules: Low Complexity Sequence Domains Form Dynamic Fibers within Hydrogels.” Cell 149 (4): 753–67.

Klopfenstein, D. V., Liangsheng Zhang, Brent S. Pedersen, Fidel Ramírez, Alex Warwick Vesztrocy, Aurélien Naldi, Christopher J. Mungall, et al. 2018. “GOATOOLS: A Python Library for Gene Ontology Analyses.” Scientific Reports 8 (1): 10872.

Krammer, Carmen, Hermann M. Schätzl, and Ina Vorberg. 2009. “Prion-like Propagation of Cytosolic Protein Aggregates: Insights from Cell Culture Models.” Prion 3 (4): 206–12.

Kraut, J. 2003. “Serine Proteases: Structure and Mechanism of Catalysis,” November. https://doi.org/10.1146/annurev.bi.46.070177.001555.

Kroschwald, Sonja, Shovamayee Maharana, Daniel Mateju, Liliana Malinovska, Elisabeth Nüske, Ina Poser, Doris Richter, and Simon Alberti. 2015. “Promiscuous Interactions and Protein Disaggregases Determine the Material State of Stress-Inducible RNP Granules.” eLife 4 (August): e06807.

Lancaster, Alex K., Andrew Nutter-Upham, Susan Lindquist, and Oliver D. King. 2014. “PLAAC: A Web and Command-Line Application to Identify Proteins with Prion-like Amino Acid Composition.” Bioinformatics 30 (17): 2501–2.

Lee, Michael. 2022. “Bit: A Multipurpose Collection of Bioinformatics Tools.” F1000Research 11 (122): 122.

Lee, Michael D. 2019. “GToTree: A User-Friendly Workflow for Phylogenomics.” Bioinformatics 35 (20): 4162–64.

Letunic, Ivica, and Peer Bork. 2007. “Interactive Tree Of Life (iTOL): An Online Tool for Phylogenetic Tree Display and Annotation.” Bioinformatics 23 (1): 127–28.

Linlin Yan. 2022. “ggvenn: Draw Venn Diagram by ‘ggplot2’”. R package version 0.1.9.

Long, John A., and Richard Cloutier. 2022. “How a 380-Million-Year-Old Fish Gave Us Fingers.” Accessed January 24, 2022. https://doi.org/10.1038/scientificamerican0620-46.

Malinovska, Liliana, Sonja Kroschwald, and Simon Alberti. 2013. “Protein Disorder, Prion Propensities, and Self-Organizing Macromolecular Collectives.” Biochimica et Biophysica Acta 1834 (5): 918–31.

Malinovska, Liliana, Sandra Palm, Kimberley Gibson, Jean-Marc Verbavatz, and Simon Alberti. 2015. “Dictyostelium Discoideum Has a Highly Q/N-Rich Proteome and Shows an Unusual Resilience to Protein Aggregation.” Proceedings of the National Academy of Sciences of the United States of America 112 (20): E2620–29.

Michelitsch, M. D., and J. S. Weissman. 2000. “A Census of Glutamine/asparagine-Rich Regions: Implications for Their Conserved Function and the Prediction of Novel Prions.” Proceedings of the National Academy of Sciences of the United States of America 97 (22): 11910–15.

Molina-García, Laura, Fátima Gasset-Rosa, María Moreno-del Álamo, M. Elena Fernández-Tresguerres, Susana Moreno-Díaz de la Espina, Rudi Lurz, and Rafael Giraldo. 2016. “Functional Amyloids as Inhibitors of Plasmid DNA Replication.” Scientific Reports 6 (1): 1–8.

Muley, Vijaykumar Yogesh, Yusuf Akhter, and Sanjeev Galande. 2019. “PDZ Domains Across the Microbial World: Molecular Link to the Proteases, Stress Response, and Protein Synthesis.” Genome Biology and Evolution. https://doi.org/10.1093/gbe/evz023.

Nan, Hao, Hongying Chen, Mick F. Tuite, and Xiaodong Xu. 2019. “A Viral Expression Factor Behaves as a Prion.” Nature Communications 10 (1): 1–11.

Nizhnikov, Anton A., Kirill S. Antonets, Stanislav A. Bondarev, Sergey G. Inge-Vechtomov, and Irina L. Derkatch. 2016. “Prions, Amyloids, and RNA: Pieces of a Puzzle.” Prion 10 (3): 182–206.

Oamen, Henry Patrick, Yasmin Lau, and Fabrice Caudron. 2020. “Prion-like Proteins as Epigenetic Devices of Stress Adaptation.” Experimental Cell Research 396 (1): 112262.

Pallarès, Irantzu, Natalia S. de Groot, Valentín Iglesias, Ricardo Sant’Anna, Arnau Biosca, Xavier Fernàndez-Busquets, and Salvador Ventura. 2018. “Discovering Putative Prion-Like Proteins in Plasmodium Falciparum: A Computational and Experimental Analysis.” Frontiers in Microbiology. https://doi.org/10.3389/fmicb.2018.01737.

Pallarès, Irantzu, and Salvador Ventura. 2017. “The Transcription Terminator Rho: A First Bacterial Prion.” Trends in Microbiology 25 (6): 434–37.

Paschkowsky, Sandra, Mehdi Hamzé, Felix Oestereich, and Lisa Marie Munter. 2016. “Alternative Processing of the Amyloid Precursor Protein Family by Rhomboid Protease RHBDL4.” The Journal of Biological Chemistry 291 (42): 21903–12.

Penalva, Ylauna Christine Megane, Sherilyn Junelle Recinto, and Lisa-Marie Munter. 2021. “Relevance of RHBDL4-Mediated APP Processing for Alzheimer’s Disease.” Alzheimer’s & Dementia: The Journal of the Alzheimer’s Association 17 Suppl 3 (December): e053878.

Price, Morgan N., Paramvir S. Dehal, and Adam P. Arkin. 2010. “FastTree 2--Approximately Maximum-Likelihood Trees for Large Alignments.” PloS One 5 (3): e9490.

Puig, Sergi, and Dennis J. Thiele. 2002. “Molecular Mechanisms of Copper Uptake and Distribution.” Current Opinion in Chemical Biology 6 (2): 171–80.

R Core Team. 2020. “R: A language and enviroment for statistical computing”. https://www.R-project.org/

Rogoza, Tatyana, Alexander Goginashvili, Sofia Rodionova, Maxim Ivanov, Olga Viktorovskaya, Alexander Rubel, Kirill Volkov, and Ludmila Mironova. 2010. “Non-Mendelian Determinant [ISP+] in Yeast Is a Nuclear-Residing Prion Form of the Global Transcriptional Regulator Sfp1.” Proceedings of the National Academy of Sciences of the United States of America 107 (23): 10573–77.

Ross, Eric D., Ulrich Baxa, and Reed B. Wickner. 2004. “Scrambled Prion Domains Form Prions and Amyloid.” Molecular and Cellular Biology 24 (16): 7206–13.

Ross, Eric D., Herman K. Edskes, Michael J. Terry, and Reed B. Wickner. 2005. “Primary Sequence Independence for Prion Formation.” Proceedings of the National Academy of Sciences of the United States of America 102 (36): 12825–30.

Ross, Eric D., Kyle S. Maclea, Charles Anderson, and Asa Ben-Hur. 2013. “A Bioinformatics Method for Identifying Q/N-Rich Prion-like Domains in Proteins.” Methods in Molecular Biology 1017: 219–28.

Rufo, Caroline M., Yurii S. Moroz, Olesia V. Moroz, Jan Stöhr, Tyler A. Smith, Xiaozhen Hu, William F. DeGrado, and Ivan V. Korendovych. 2014. “Short Peptides Self-Assemble to Produce Catalytic Amyloids.” Nature Chemistry 6 (4): 303–9.

Sabaté, Raimon, Montserrat Gallardo, and Joan Estelrich. 2003. “An Autocatalytic Reaction as a Model for the Kinetics of the Aggregation of β-Amyloid.” Biopolymers. https://doi.org/10.1002/bip.10441.

Sabate, Raimon, Frederic Rousseau, Joost Schymkowitz, and Salvador Ventura. 2015. “What Makes a Protein Sequence a Prion?” PLoS Computational Biology 11 (1): e1004013.

Shen, Wei, and Hong Ren. 2021. “TaxonKit: A Practical and Efficient NCBI Taxonomy Toolkit.” Journal of Genetics and Genomics = Yi Chuan Xue Bao 48 (9): 844–50.

Shi, Yanfang, Xiaohui Li, Guangzhu Ding, Yangjiang Wu, Yuyan Weng, and Zhijun Hu. 2014. “Control of β-Sheet Crystal Orientation and Elastic Modulus in Silk Protein by Nanoconfinement.” Macromolecules. https://doi.org/10.1021/ma501864g.

Slotta, Ute, Simone Hess, Kristina Spiess, Thusnelda Stromer, Louise Serpell, and Thomas Scheibel. 2007. “Spider Silk and Amyloid Fibrils: A Structural Comparison.” Macromolecular Bioscience 7 (2): 183–88.

Su, Ting-Yi, and Paul M. Harrison. 2019. “Conservation of Prion-Like Composition and Sequence in Prion-Formers and Prion-Like Proteins of Saccharomyces Cerevisiae.” Frontiers in Molecular Biosciences. https://doi.org/10.3389/fmolb.2019.00054.

Tetz, George, and Victor Tetz. 2017. “Prion-Like Domains in Phagobiota.” Frontiers in Microbiology 8 (November): 2239.

Tetz, George, and Victor Tetz.. 2018. “Prion-like Domains in Eukaryotic Viruses.” Scientific Reports 8 (1): 8931.

UniProt Consortium. 2021. “UniProt: The Universal Protein Knowledgebase in 2021.” Nucleic Acids Research 49 (D1): D480–89.

Urban, Sinisa, and Seth W. Dickey. 2011. “The Rhomboid Protease Family: A Decade of Progress on Function and Mechanism.” Genome Biology. https://doi.org/10.1186/gb-2011-12-10-231.

Urban, Siniša, and Syed M. Moin. 2014. “A Subset of Membrane-Altering Agents and γ-Secretase Modulators Provoke Nonsubstrate Cleavage by Rhomboid Proteases.” Cell Reports 8 (5): 1241–47.

Vallota-Eastman, Alec, Eleanor C. Arrington, Siobhan Meeken, Simon Roux, Krishna Dasari, Sydney Rosen, Jeff F. Miller, David L. Valentine, and Blair G. Paul. 2020. “Role of Diversity-Generating Retroelements for Regulatory Pathway Tuning in Cyanobacteria.” BMC Genomics 21 (1): 664.

Volkov, Kirill V., Anna Yu Aksenova, Malle J. Soom, Kirill V. Osipov, Anton V. Svitin, Cornelia Kurischko, Irina S. Shkundina, Michael D. Ter-Avanesyan, Sergey G. Inge-Vechtomov, and Ludmila N. Mironova. 2002. “Novel Non-Mendelian Determinant Involved in the Control of Translation Accuracy in Saccharomyces Cerevisiae.” Genetics 160 (1): 25–36.

Wang, N., S. Gottesman, M. C. Willingham, M. M. Gottesman, and M. R. Maurizi. 1993. “A Human Mitochondrial ATP-Dependent Protease That Is Highly Homologous to Bacterial Lon Protease.” Proceedings of the National Academy of Sciences of the United States of America 90 (23): 11247–51.

Westaway, David, Nathalie Daude, Serene Wohlgemuth, and Paul Harrison. 2011. “The PrP-like Proteins Shadoo and Doppel.” Topics in Current Chemistry 305: 225–56.

White, Mark J., Hongjun He, Renee M. Penoske, Sally S. Twining, and Thomas C. Zahrt. 2010. “PepD Participates in the Mycobacterial Stress Response Mediated through MprAB and SigE.” Journal of Bacteriology 192 (6): 1498–1510.

Wickham, Hadley. ggplot2: elegant graphics for data analysis. springer, 2016.

Wickner, R. B. 1994. “[URE3] as an Altered URE2 Protein: Evidence for a Prion Analog in Saccharomyces Cerevisiae.” Science.

Wickner, Reed B., Herman K. Edskes, and Frank Shewmaker. 2006. “How to Find a Prion: [URE3], [PSI+] and [beta].” Methods 39 (1): 3–8.

Wickner, Reed B., and Amy C. Kelly. 2016. “Prions Are Affected by Evolution at Two Levels.” Cellular and Molecular Life Sciences: CMLS 73 (6): 1131–44.

Yoshimura, Yuichi, Yuxi Lin, Hisashi Yagi, Young-Ho Lee, Hiroki Kitayama, Kazumasa Sakurai, Masatomo So, Hirotsugu Ogi, Hironobu Naiki, and Yuji Goto. 2012. “Distinguishing Crystal-like Amyloid Fibrils and Glass-like Amorphous Aggregates from Their Kinetics of Formation.” Proceedings of the National Academy of Sciences of the United States of America 109 (36): 14446–51.

Yuan, Andy H., and Ann Hochschild. 2017. “A Bacterial Global Regulator Forms a Prion.” Science 355 (6321): 198–201.

Zajkowski, Tomasz, Michael D. Lee, Shamba S. Mondal, Amanda Carbajal, Robert Dec, Patrick D. Brennock, Radoslaw W. Piast, et al. 2021. “The Hunt for Ancient Prions: Archaeal Prion-Like Domains Form Amyloid-Based Epigenetic Elements.” Molecular Biology and Evolution 38 (5): 2088–2103.

Zambrano, Rafael, Oscar Conchillo-Sole, Valentin Iglesias, Ricard Illa, Frederic Rousseau, Joost Schymkowitz, Raimon Sabate, Xavier Daura, and Salvador Ventura. 2015. “PrionW: A Server to Identify Proteins Containing Glutamine/asparagine Rich Prion-like Domains and Their Amyloid Cores.” Nucleic Acids Research 43 (W1): W331–37.

Zaremba-Niedzwiedzka, Katarzyna, Eva F. Caceres, Jimmy H. Saw, Disa Bäckström, Lina Juzokaite, Emmelien Vancaester, Kiley W. Seitz, et al. 2017. “Asgard Archaea Illuminate the Origin of Eukaryotic Cellular Complexity.” Nature 541 (7637): 353–58.

