## Supplemental materials list for "Universal functions of prion candidates across all three domains of life suggest a primeval role of protein self-templating"

### Supplemental Figures

Figure S1: Same trees as main figures 1, 2, and 3, but overlaid in blue here are those with candidate prions that were annotated with the calcium ion binding (GO:0005509) GO term.

Figure S2: Same trees as main figures 1, 2, and 3, but overlaid in blue here are those with candidate prions that were annotated with the helicase activity (GO:0004386) GO term.

### Supplemental Tables

Table S1: Reference proteomes info

Table S2: Proteome-level PLAAC COREScore >0 summaries

Table S3: Archaea GO enrichment analysis results

Table S4: Bacteria GO enrichment analysis results

Table S5: Eukarya GO enrichment analysis results

Table S6: KO annotations of proteins associated with GO:0016567

Table S7: PLAAC-positive protein counts grouped by GO term and taxonomic domain

Table S8: GO terms enriched in both Bacteria and Archaea

Table S9: PLAAC results and sequences of proteins with enriched GO terms found in all 3 domains

Table S10: Enriched GO terms with their corresponding proteins’ KO annotations

Table S11: PLAAC results and sequences of proteins with enriched GO terms found in Bacteria and Archaea

Table S12: Summary table of discussed GO terms and their corresponding proteins’ KO annotations
