## Supplemental Figure 2 for "Universal functions of prion candidates across all three domains of life suggest a primeval role of protein self-templating"

Figure S2 | Same trees as main figures 1, 2, and 3, but overlaid in blue here are those with candidate prions that were annotated with the helicase activity (GO:0004386) GO term.

### Archaea

Tree scale: 1.0

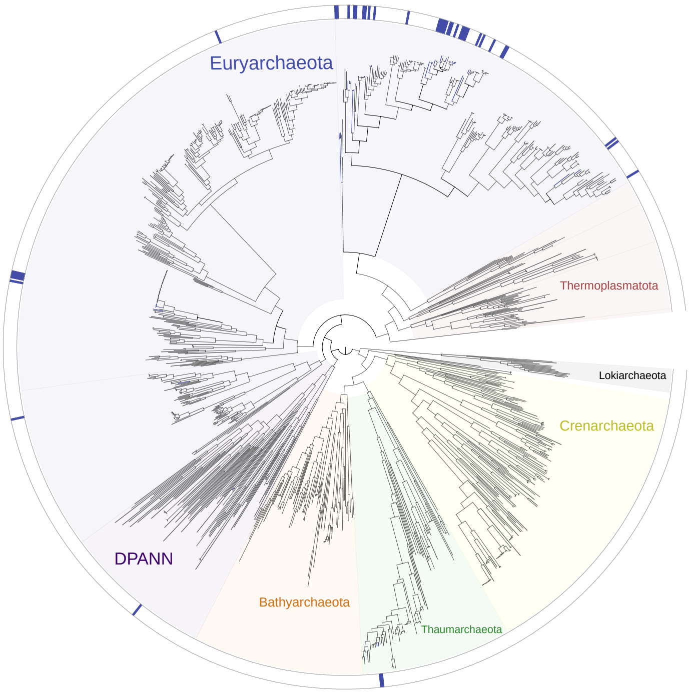


### Bacteria

Tree scale: 1.0

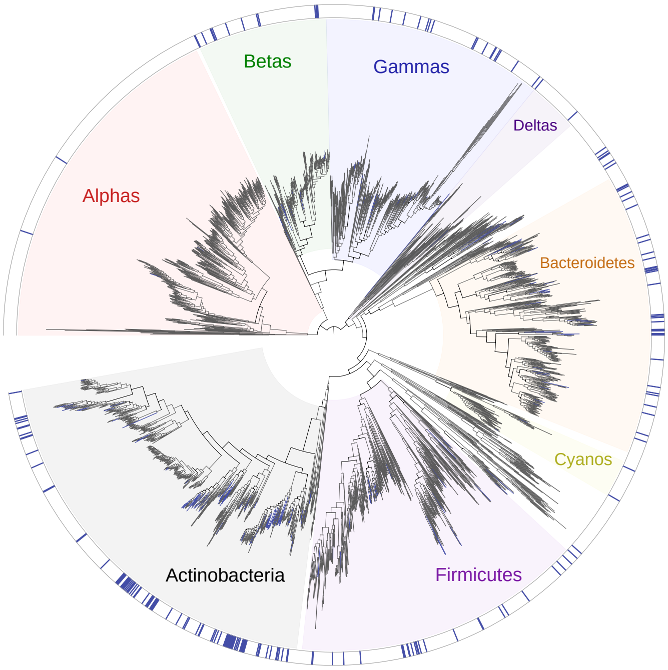


### Eukarya

Tree scale: 1.0

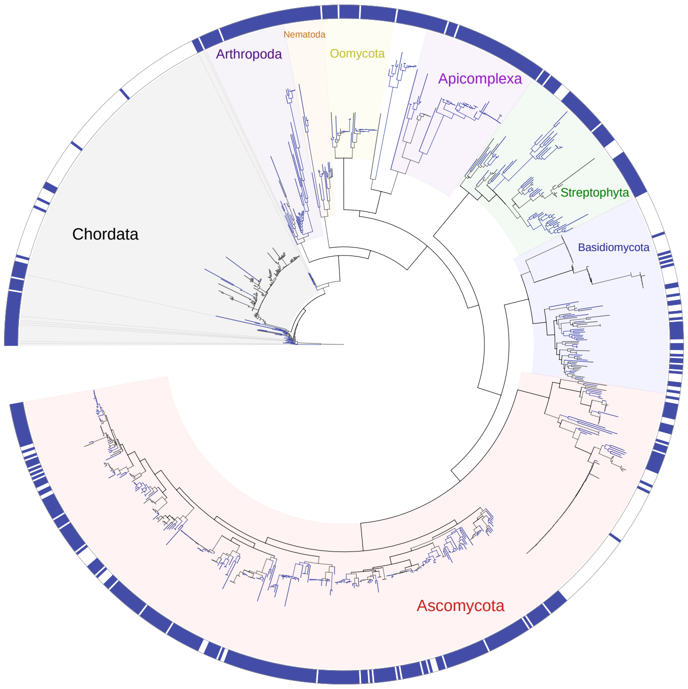
